# Whole-transcriptome-scale and isoform-resolved spatial imaging of single cells in complex tissues

**DOI:** 10.1101/2025.08.27.672533

**Authors:** Limor Cohen, Aaron Halpern, Timothy R. Blosser, Z. Phil Che, Xingjie Pan, Xiaowei Zhuang

## Abstract

Cell and tissue functions arise from complex interactions among numerous genes, and a systematic understanding of these functions requires isoform-resolved, whole-transcriptome-scale analysis of single cells with high spatial resolution. Here, we introduce a spatial transcriptomics method based on a novel *in situ* RNA amplification strategy, enabling short RNA sequence detection and hence spatially resolved expression profiling of individual cells at the whole-transcriptome scale with splice-isoform resolution. Using this approach, we imaged ∼33,000 distinct RNAs—including ∼23,000 genes and ∼10,000 isoform-defining transcripts—in the mouse brain. Our data enabled systematic analyses of region- and cell-type-specific gene programs and ligand-receptor-based cell-cell communications. These data further revealed a rich spatial diversity and cell-type specificity in isoform usage across numerous genes and identified brain structures particularly enriched for specific isoform usage. We anticipate broad application of this method for molecular and cellular analysis of tissues, unlocking previously inaccessible discoveries in cell and organismal biology.

## INTRODUCTION

Recent advances in spatial transcriptomics technologies have enabled genome-scale profiling of RNA expression in intact tissues, which allows *in situ* identification and spatial mapping of molecularly distinct cell types and associated gene-regulatory mechanisms, providing rich insights into the molecular and cellular architecture of complex tissues.^1–4^ Two major classes of spatial transcriptomics methods have emerged based on multiplexed imaging and next-generation sequencing.^1, 2, 4^ However, these approaches face a major tradeoff between spatial resolution and transcriptomic coverage. Imaging-based methods provide high spatial resolution, routinely achieving subcellular resolution, but are typically limited to detecting a targeted set of gene. In contrast, sequencing-based methods offer untargeted coverage of the whole genome, but have limited spatial resolution, making it challenging to achieve single-cell or subcellular resolution.

The ability to perform transcriptome-wide RNA profiling with high spatial resolution is therefore in critical demand. At single-cell resolution, such measurements would enable unbiased identification of cell types and states and comprehensive analysis of gene programs, revealing both global organization and local microenvironment of cells, as well as how gene expression is regulated across cell types, states, and spatial neighborhoods. Transcriptome-wide RNA measurements with subcellular resolution can further provide insights into post-transcriptional regulatory mechanisms—such as local translation and RNA degradation—that shape the spatiotemporal dynamics of protein expression. Moreover, many genes have different isoforms of transcripts due to alternative splicing, differential transcription start site (TSS) usage, and other forms of variation.^5^ This isoform diversity underlies critical aspects of cell identity, signaling, and disease, ^6–8^ yet current spatial transcriptomics methods—both imaging- and sequencing-based approaches—lack the ability to systematically resolve isoforms at the genome scale with single-cell resolution. Thus, how different isoforms are differentially expressed spatially and across cell types in tissues remains poorly understood for most genes. Transcriptome-wide spatial profiling of single cells would fill this major gap.

Some state-of-the-art sequencing-based approaches provide spatially resolved, untargeted transcriptome measurements at single-cell or near single-cell resolution, but have relatively limited cell or transcript capture efficiency.^9–12^ The high sequencing cost could also hinder the wide adoption of these approaches for organ-level characterization, which often requires profiling of hundreds of thousands to millions of cells. Furthermore, high-throughput sequencing employed in these methods typically probes only the 3’ and/or 5’ ends of the transcripts, hence missing most isoform information. While recent development of spatial long-read sequencing can recover full-length transcripts,^13–15^ these approaches currently have limited spatial resolution, falling short of resolving single cells.

Imaging-based methods provides single-cell and subcellular resolution, but have yet to achieve whole-transcriptome coverage. For example, multiplexed FISH has achieved simultaneous imaging of thousands of genes in individual cells and allows flexible coding schemes to avoid signal crowding by adjusting the fraction of RNAs imaged per imaging round.^16–19^ However, precise RNA identification typically requires tens of FISH probes per transcript, preventing detection of short RNA transcripts or transcripts differing by short sequences, such as splice isoforms. In addition, the high probe cost per gene library for whole-transcriptome measurements limits scalability and wide adoption. *In situ* sequencing^20–22^ can detect short transcripts, but detects a fixed fraction (∼25%) of genes per imaging round, leading to a signal crowding problem arising from the high transcript density, which has also prevented transcriptome-wide measurement.

RNA signal amplification offers a potential solution by reducing the number of probes needed per transcript in multiplexed FISH, thereby relaxing transcript-length constraints and lowering probe cost. Several amplification methods have been reported, including rolling circle amplification (RCA),^23–25^ branched DNA amplification,^26, 27^ hybridization chain reaction,^28, 29^ ClampFISH,^30^ and signal amplification by exchange reaction,^31^ and some have been integrated with spatial transcritomics.^20–22, 32–35^ Nonetheless, isoform-resolved, whole-transcriptome-scale imaging has not yet been achieved.

To overcome these limitations and enable whole-transcriptome-scale and isoform-resolved imaging, we developed an *in-situ* RNA amplification method, termed RT&T-AMP, which uses reverse transcription followed by transcription to generate RNA amplicons at endogenous transcript sites, and we further integrated RT&T-AMP with multiplexed error-robust fluorescence in situ hybridization (MERFISH)^16^ for single-cell transcriptome imaging. Using this approach, we demonstrated whole-transcriptome-scale imaging of ∼23,000 genes—including both protein-coding and non-coding RNAs—in single cells within mouse brain tissue sections. Our approach allowed systematic analyses of (i) differential gene expression across brain regions and microenvironments, (ii) spatially correlated gene modules that align with or define anatomic structures, and (iii) region- and cell-type-specific cell-cell signaling interactions. Furthermore, we demonstrated simultaneous imaging of 33,000 distinct RNAs, including ∼23,000 genes and 10,000 isoforms. Our data revealed numerous genes with region- and cell-type-specific isoform usage in the brain. We additionally identified brain structures that are particularly enriched for specific isoform usage across many genes. Together, these results establish a powerful and scalable technology for spatial transcriptomic profiling of single cells at the whole-transcriptome scale with isoform resolution.

## RESULTS

### *In-situ* RNA amplification with reverse transcription and transcription (RT&T-AMP)

To overcome the transcript-length constraints and increase the signal-to-noise ratio in single-cell transcriptome imaging, we developed RT&T-AMP, which amplifies signals from RNA molecules *in situ* using reverse transcription (RT) followed by transcription, generating RNA amplicons localized to the original transcript sites (**Figure 1A**). First, we performed RT *in situ* using a poly-T primer that bound to the poly-A tail of the cellular RNA, producing a complementary DNA (cDNA) with a short non-templated CCC extension at the 3′ end. This CCC overhang was then hybridized to a template switching oligo (TSO) containing the T7 promoter sequence, allowing the reverse transcriptase to switch templates from the cellular RNA to the TSO and continue extending the cDNA. The resulting cDNA thus carried a T7 promoter at its 3′ end. We then added T7 RNA polymerase to the tissue sample to transcribe each cDNA into multiple copies of RNA, generating an RNA amplicon at the location of each original transcript.

**Figure 1.**
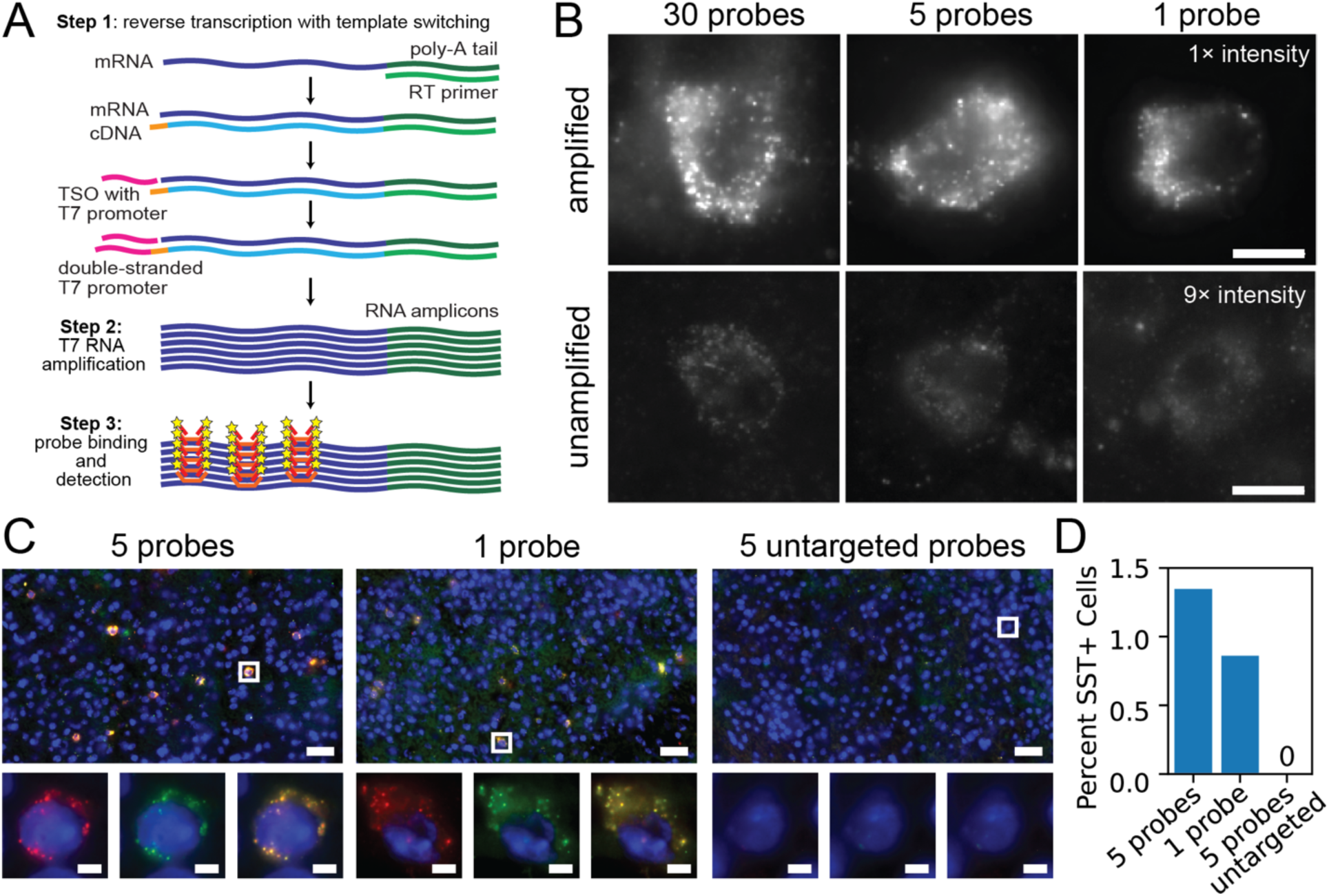
*In situ* RNA amplification by reverse transcription and transcription (RT&T-AMP) (A) Schematic of the RT&T-AMP workflow. Reverse transcription with a polyT primer generates a complementary DNA (cDNA) and adds a short non-templated CCC overhang to the cDNA. A template switching oligo (TSO) containing a T7 promoter sequence and GGG to bind to the CCC overhand allows template switching of the reverse transcriptase to add the T7 promoter to the cDNA. T7 RNA polymerase then transcribes the cDNA into multiple copies of RNA, forming an amplicon. MERFISH encoding probes and fluorescently labeled readout probes are then used to detect the RNA amplicons. (B) smFISH images of an inhibitory neuronal subtype marker *Sst* in single cells in the mouse brain tissue using probe sets containing 30, 5, and 1 encoding probes with (top) and without (bottom) RT&T-AMP. Scale bars: 10 µm. (C) Left and middle: Two-color smFISH images of *Sst* from multiple fields of view (FOV) tiled together, showing high co-localization of signals from probes labeled with different fluorescent dyes. Left: detection using 5 encoding probes per gene. Middle: detection using 1 encoding probe per gene. Since each amplicon contains multiple copies of RNA, multiple probes can bind to the same amplicon, allowing colocalization of the differently colored signals. Stray, non-specific binding of probes is unlikely to generate colocalized signal. Right: smFISH image with non-targeting probes (5 probes per gene) yield negligible signal. Bottom panels show the amplified view of the boxed region in the top panels. Scale bars: 10 µm (top), 5 µm (bottom). (D) Percent of cells in the imaged brain sections showing positive *Sst* signal, detected using 5 encoding probes, 1 encoding probes, and 5 non-targeting probes.

To validate RT&T-AMP, we performed single-molecule FISH (smFISH)^36, 37^ experiments targeting a cell-type-specific gene, *Sst*, with 30, 5, or 1 probe(s), each containing 30-nt target-binding sequence. RT&T-AMP produced drastically brighter signals for RNA molecules than unamplified smFISH (**Figure 1B**). While RT&T-AMP enabled robust detection of individual RNA molecules even with only a single probe, unamplified smFISH generated weak or undetectable signals when a small number of probes (1 or 5) were used. In addition, we performed two-color smFISH of *Sst* with probes labeled by different fluorophores and observed nearly perfect colocalization between the differently colored probes (**Figure 1C**, **left and middle panels**), demonstrating high detection specificity. As expected for *Sst*, which is expressed in a specific inhibitory neuronal subtype, only a small subset of cells showed positive smFISH signal (**Figure 1C, left and middle panels**, and **Figure 1D**). Moreover, control experiments with non-targeting FISH probes generated essentially no detectable signal in any cell (**Figure 1C, right panel**, and **Figure 1D**), confirming the high specificity of RT&T-AMP-enabled smFISH with a small number of probes.

### Whole-transcriptome-scale MERFISH with RT&T-AMP in brain tissues

We next integrated RT&T-AMP with MERFISH to perform whole-transcriptome-scale imaging in mouse brain tissue, targeting ∼23,000 genes with six probes per gene (**Figure 2A**). To minimize signal crowding, we split the gene panel into two groups and imaged them with two separate, back-to-back MERFISH runs on the same sample; each group was imaged using an error-correcting, 90-bit Hamming-distance-4 and Hamming-weight-4 code. Imaging both groups on the same sample required 180 bits in total, which were read out using 60 rounds of three-color imaging.

**Figure 2.**
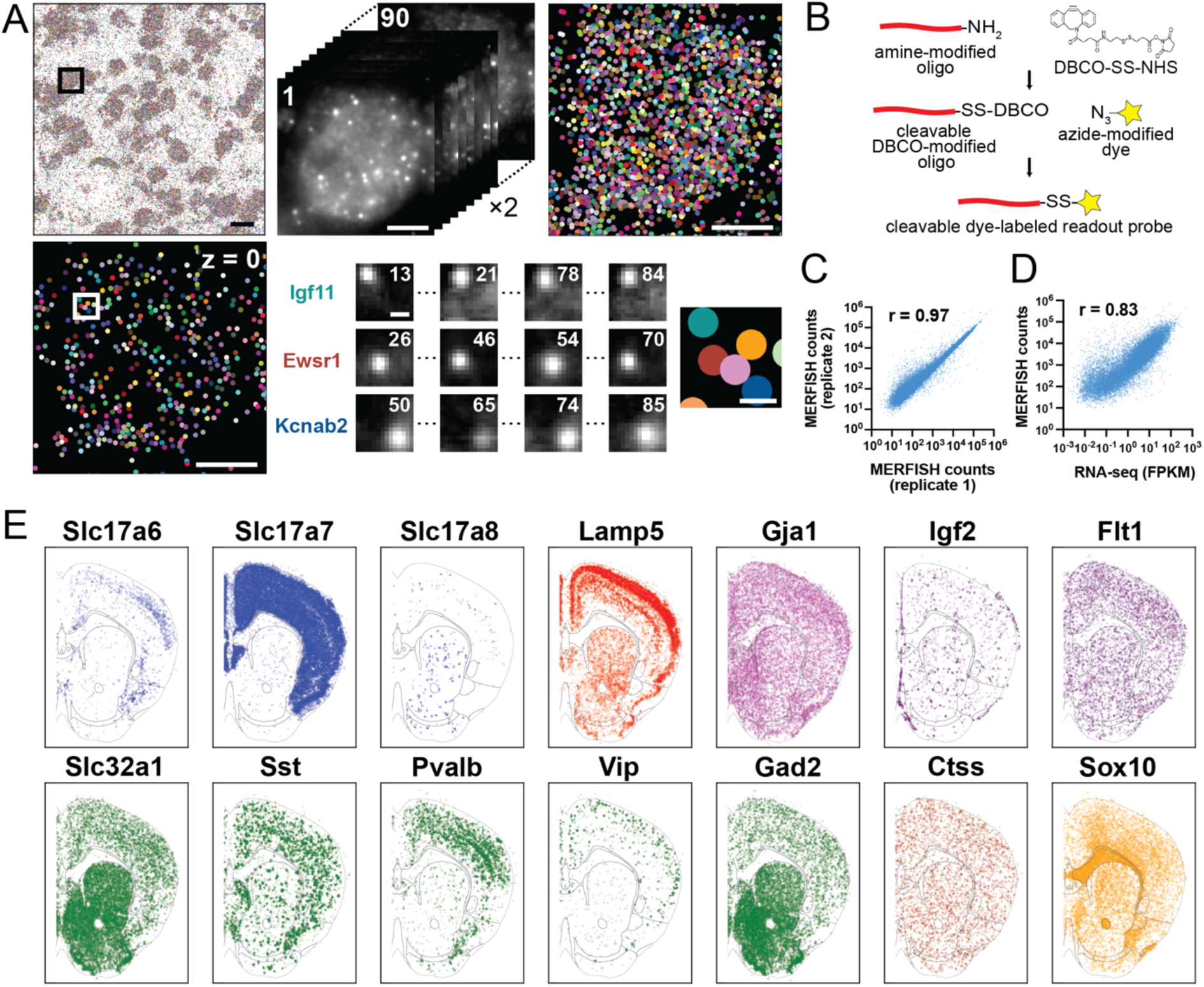
Whole-transcriptome-scale MERFISH with RT&T-AMP in mouse brain tissues. (A) Top left: RT&T-AMP-MERFISH image for ∼23,000 genes in one field of view, with molecules colored by their genetic identity. Top middle and right: Raw, per-bit images and decoded MERFISH image of a single cell (projection image across multiple z planes) corresponding to the boxed region in the left panel. Bottom left: image of decoded RNA molecules in one z plane for the same cell. Bottom middle: Raw, per-bit images of a small area corresponding to the boxed region in the left panel, showing the four on-bit images for each of the three indicated genes. Bottom right: decoded RNAs colored by genetic identity in the small area. Scale bars: 20 µm (top left), 5 µm (top middle, top right, bottom left), 500 nm (bottom middle, bottom right). (B) Workflow to generate the cleavable, dye-labeled readout probes via DBCO–azide click chemistry. (C) Scatter plot of RNA counts for the ∼23,000 genes showing high reproducibility of whole-transcriptome-scale RT&T-AMP-MERFISH between replicates. (D) Scatter plot showing high correlation between RNA counts of the ∼23,000 genes determined by RT&T-AMP-MERFISH and the average expression level (in FPKM) of the corresponding genes determined by bulk RNA-seq. (E) Spatial maps of representative marker genes for major cell classes, showing expected spatial distributions in a representative coronal brain section. The grey lines demarcate brain regions defined in the Allen CCF.

MERFISH barcoding and decoding was implemented by hybridizing encoding probes to the target RNAs, each probe containing a target-binding sequence and multiple readout sequences^16^. The collection of readout sequences binding to each RNA determines the binary barcode of the gene, with each present readout sequence indicating “1” for the corresponding bit in the barcode. We then detected the readout sequences by sequential rounds of hybridization with dye-labeled, complementary readout probes. To enable fast signal removal between hybridization rounds, dyes were linked to the readout probe via a di-sulfide bond that can be cleaved by TECP.^38^

A 180-bit MERFISH measurement requires 180 unique readout probes, which would cost >$100,000 when acquired from commercial sources. To reduce cost, we developed a cost-effective protocol for generating such cleavable, dye-labeled readout probes through click chemistry (**Figure 2B**) at a fraction of the commercial cost (∼$20 per probe).

Using RT&T-AMP to amplify the endogenous RNAs, 6 encoding probes to target each gene, and the afore-mentioned custom-made readout probes, we performed MERFISH imaging of the ∼23,000 genes in coronal sections of the mouse brain. We imaged four brain slices, with two slices located in an anterior region and two slices located in a posterior region (∼150,000 cells in total). Single RNA molecules were clearly detected and decoded in individual cells (**Figure 2A**). Our RT&T-AMP-MERFISH data were highly reproducible between replicates (**Figure 2C**) and the average expression levels of individual genes measured by RT&T-AMP-MERFISH showed excellent correlation with bulk RNA sequencing results (**Figure 2D**).

In addition to enabling measurements of RNAs too short to be detected without amplification, RT&T-AMP also greatly reduced the probe cost of MERFISH imaging experiments. Without amplification, each gene would typically require ∼50 targeting probes and imaging of ∼23,000 genes would require >1.1 million encoding probes in total, costing ∼$50,000-$100,000. Here, with RT&T-AMP, we used 6 probes per gene and our smFISH experiments indicated the feasibility of using fewer probes (**Figure 1B-D**). Thus, RT&T-AMP reduced probe requirement and cost by ∼10 fold or more. Combined with the reduced readout probe cost as described above, this makes whole-transcriptome imaging experiments highly affordable. Moreover, the drastic increase in RNA signal by RT&T-AMP also offers the possibility of using low N.A. objectives with a greater field of view, increasing the imaging throughput (number of cells imaged per unit time).

### Spatial distributions and spatial modules of genes

To facilitate spatial analysis of gene expression, we aligned our MERFISH images to the Allen Mouse Brain Common Coordinate Framework (CCF, version 3)^39^ (**Figure S1**). Visual inspection of marker genes for major cell types, including excitatory neurons (e.g. *Slc17a7*, *Slc17a6*, *Slc17a8*), inhibitory neurons (e.g. *Slc32a1*, *Gad2*, *Lamp5*, *Pvalb*, *Sst*, *Vip*), astrocytes (e.g. *Gja1*), oligodendrocytes (e.g. *Sox10*), microglia/immune cells (e.g. *Ctss*), and vascular and ventricular cells (e.g. *Flt 1* and *Ifg2*), showed expected spatial distributions (**Figure 2E**), validating our measurements. For example, excitatory neuronal markers, *Slc17a7* (Vglut1), *Slc17a6* (Vglut2) and *Slc17a8* (Vglut3), were enriched in different brain regions with *Slc17a7* highly enriched in the isocortex, cortical subplate and olfactory areas; inhibitory neuronal markers (*Slc32a1* and *Gad2*) were highly enriched in the striatum, which is dominated by inhibitory neurons; specific inhibitory neuronal subtype markers, *Lamp5* (which also marks upper-layer excitatory neurons), *Vip*, *Sst*, and *Pvalb*, showed different enrichment in upper and lower cortical layers; the oligodendrocyte marker, *Sox10*, was enriched in the fiber tract; the *VLMC* marker, *Ifg2*, was enriched at the meninges – all consistent with prior knowledge. ^40, 41^

To systematically investigate the spatial distributions of the ∼23,000 genes and their correlation with each other, we performed spatial Hotspot analysis ^42^ on the imaged coronal sections (**Figure 3; Figures S2 and S3; Table S3**). In this analysis, the correlation of spatial expression patterns was computed between each pair of genes and groups of genes with spatially correlated expression were identified as gene modules (**Methods**).

**Figure 3.**
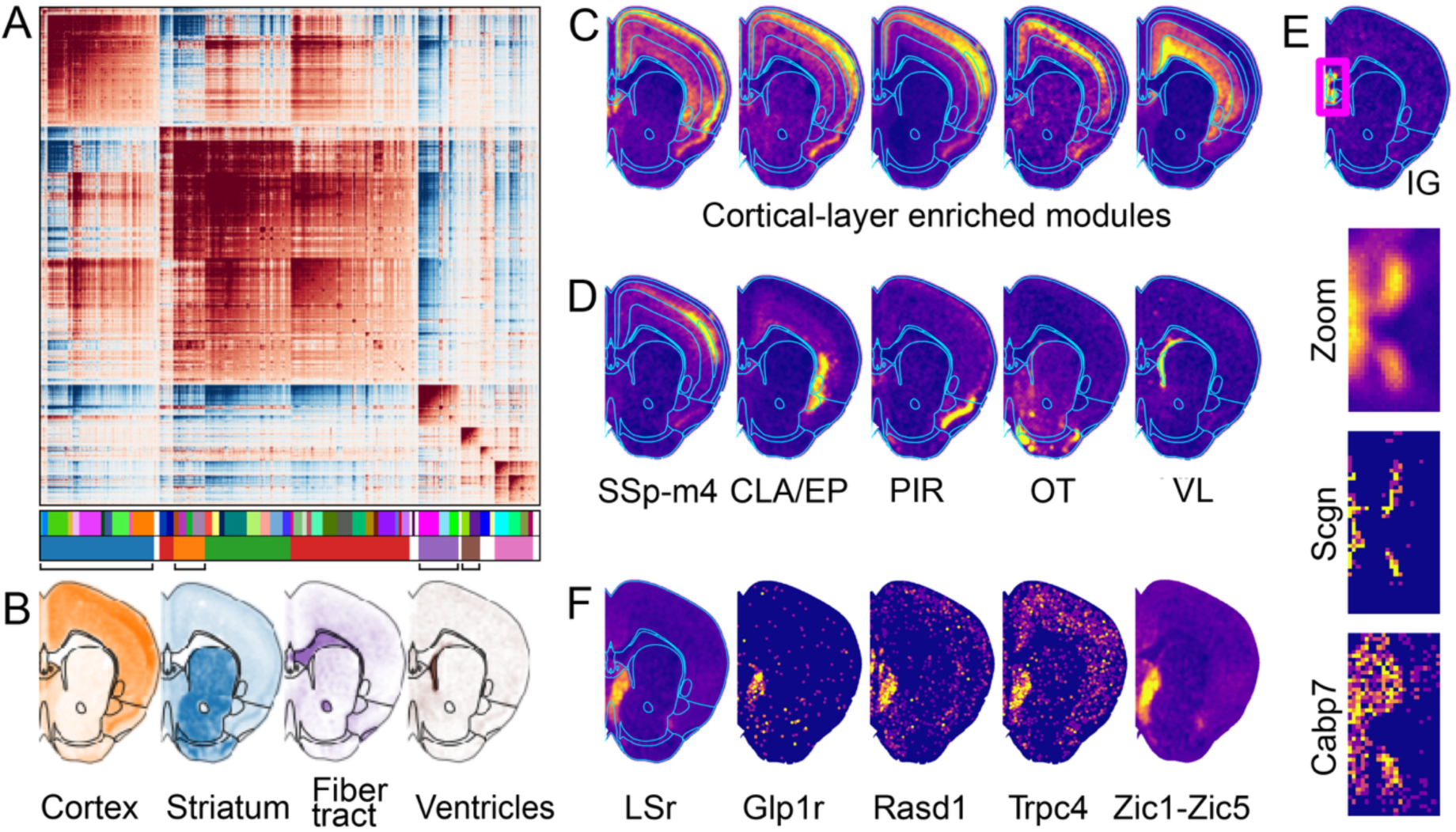
Spatial Hotspot analysis identifies gene modules aligned with anatomical structures in the brain. (A) Hotspot analysis of the whole-transcriptome-scale RT&T-AMP-MERFISH data to identify group of genes with spatially correlated expression, defining modules that align with anatomical structures at multiple resolutions. Spatial correlation matrix and hierarchical clustering generate gene modules at a coarse resolution (≥300 genes per module, outer bar) and a finer resolution (≥30 genes per module, inner bar). Results for the two anterior sections are shown in this figure and Figure S2; results for the two posterior sections are shown in Figure S3. (B) Example coarse-resolution gene modules that align with several major anatomical regions and structures. The black lines in (B) and blue lines in (C-F) demarcate brain regions defined in the Allen CCF. (C) Example fine-resolution gene modules that show enrichment in different cortical layers. (D) Example fine-resolution modules corresponding to small brain structures, including the primary somatosensory area, layer 4 (SSp-m4), the claustrum (CLA) and endopiriform nucleus (EP), the piriform area (PIR), the olfactory tubercle (OT), and the lateral ventricle (VL). (E) The indusium griseum (IG) gene module, containing *Scgn* and *Cabp7*, among other genes (33 genes in total). (F) The LSr gene module, containing *Glp1r*, *Rasd1*, *Trpc4*, and *Zic* transcription factors, among other genes (207 genes in total).

The two adjacent anterior slices showed highly similar results (**Figure S2A, B**). Of the ∼23,000 genes, Hotspot analysis found ∼10,000 genes that exhibited spatially non-random expression patterns and identified gene modules at different resolutions by varying the minimum gene number per module (**Figure 3A**; **Figure S2C**). At a coarse resolution (gene number per module threshold = 300), we identified seven modules, which were aligned with major anatomical regions, including the isocortex, cortical subplate and olfactory areas (Module A-II), striatum (Module A-I), fiber tracts (Module A-V), ventricles (Module A-VI), and meninges (Module A-VII) (**Figure 3B; Figure S2C**). Two large modules (Modules A-III and A-IV) spanned the entire cerebrum, but were distinct in the correlation matrix, suggesting that they may comprise finer modules at a higher resolution. Indeed, higher-resolution analysis (gene number per module threshold = 30) revealed ∼50 finer gene modules (**Figure 3A**; **Figure S2C**) that subdivided the low-resolution modules and aligned with smaller brain regions. For example, a number of layer-like modules were resolved in the cortex (**Figure 3C**; **Figure S2C**). Many other modules corresponding to small brain regions or anatomical structures were also observed (**Figure 3D, E**; **Figure S2C**). As an example, the Indusium Griseum (IG) module (**Figure 3E**) contained 33 genes, including not only known IG markers, such as *Scgn*, but also many other genes, such as *Cabp7*, which is consistent with the IG’s proposed molecular similarity to the CA2 region of the hippocampus.^43^ The two posterior slices, spaced ∼50 μm apart, showed partially overlapping but distinct modules due to rapid anatomical transitions along the anterior-posterior axis in this region (**Figure S3**), as observed in the Allen CCF.^39^ As in the anterior slices, many high-resolution modules were identified, which aligned with specific brain regions, such as different cortical layers, different hippocampal regions including the dentate gyrus (DG), CA1, and CA2/3, as well as many other small nuclei, such as the parafascicular nucleus (P1-I-2) (**Figure S3**).

As a specific interesting example, we describe the LSr module in the anterior slices here. This module consisted of 207 genes localized to the lateral septal nucleus, rostral part (LSr) of the striatum (**Figure 3F**). This module included the Zic family of transcription factors, consistent with previous knowledge of the expression of these genes in the LSr.^44, 45^ Interestingly, we also observed spatial colocalization of *Glp1r*, a G protein-coupled receptor (GPCR) gene that regulates blood glucose levels,^46, 47^ and *Trpc4*, a calcium channel gene in downstream pathways of *Glp1r*,^48^ in the LSr module. Since *Trpc4* and *Glp1r* have been implicated in neural circuits underlying social interactions,^49–51^ and the LSr is a brain region involved in emotion and social behaviors, ^52, 53^ our data suggest the possibility that *Glp1r* and *Trpc4* signaling contributes to the role of LSr in the regulation of social behaviors. Another interesting gene in this module is *Rasd1*, a guanine nucleotide exchange factor gene involved in G-protein signaling.^54^ Both *Rasd1* and *Glp1r* are known to influence cAMP signaling but in opposite directions: *Rasd1* inhibits adenylyl cyclase, reducing cAMP production, whereas *Glp1r* activates adenylyl cyclase, increasing cAMP production.^54^ cAMP is a key second messenger in stress and emotion-related signaling. ^52, 55^ Our observed co-expression of *Rasd1* and *Glp1r* in the LSr thus suggests a mechanism by which opposing regulation of cAMP by *Rasd1* and *Glp1r* could shape and modulate LSr’s contribution to emotion and behavioral control. These results illustrate the power of whole-transcriptome-scale spatial analysis to generate interesting hypotheses to be tested by future functional studies.

### Cell-type-specific differential gene expression across different brain regions

Next, to facilitate cell-type-specific spatial analysis of gene expression, we performed *de novo* clustering of cells based on their gene expression profiles and identified known major cell classes and subtypes of neurons. Major cell classes included neurons, astrocytes, mature oligodendrocytes, newly formed oligodendrocytes (NFOLs), oligodendrocyte precursor cells (OPCs), microglia, ependymal cells, cells of the choroid plexus, endothelial cells, pericytes, red blood cells (RBCs), smooth muscle cells (SMCs), and vascular leptomeningeal cells (VLMCs) (**Figure 4A**; **Figure S4A**). Neuronal subtypes included cortical excitatory neurons of different projection types and cortical layers (L2/3 intratelencephalic (IT), L4/5 IT, L5 IT, L6 IT, L2/3 IT RSP, L4/5 IT RSP, L5 extratelencephalic (ET), L5 near-projecting (NP), L6 corticothalamic (CT), and L6b),^56, 57^ inhibitory neurons marked by canonical markers *Pvalb*, *Sst*, *Vip*, and *Lamp5*, hippocampal neurons localized to the CA1, CA2/3, DG, and IG, two types of medium spiny neurons (MSNs, D1- and D2-type) localized to striatum, as well as other neuronal subtypes localized to striatum, cortical subplate, olfactory areas, thalamus, hypothalamus, and midbrain (**Figure 4B**; **Figure S4B, C**). The *de novo* clustering results were in agreement with classification results obtained from scRNA-seq label transfer based on integration of our RT&T-AMP-MERFISH data with previous scRNA-seq data^40^ (**Figure S4D**). These transcriptome-based cell-type classification results were also consistent with the known spatial locations of these cell types. Of note, we classified cells at a medium level of granularity, mostly at the subclass level, because our subsequent analyses were only performed at this level.

**Figure 4.**
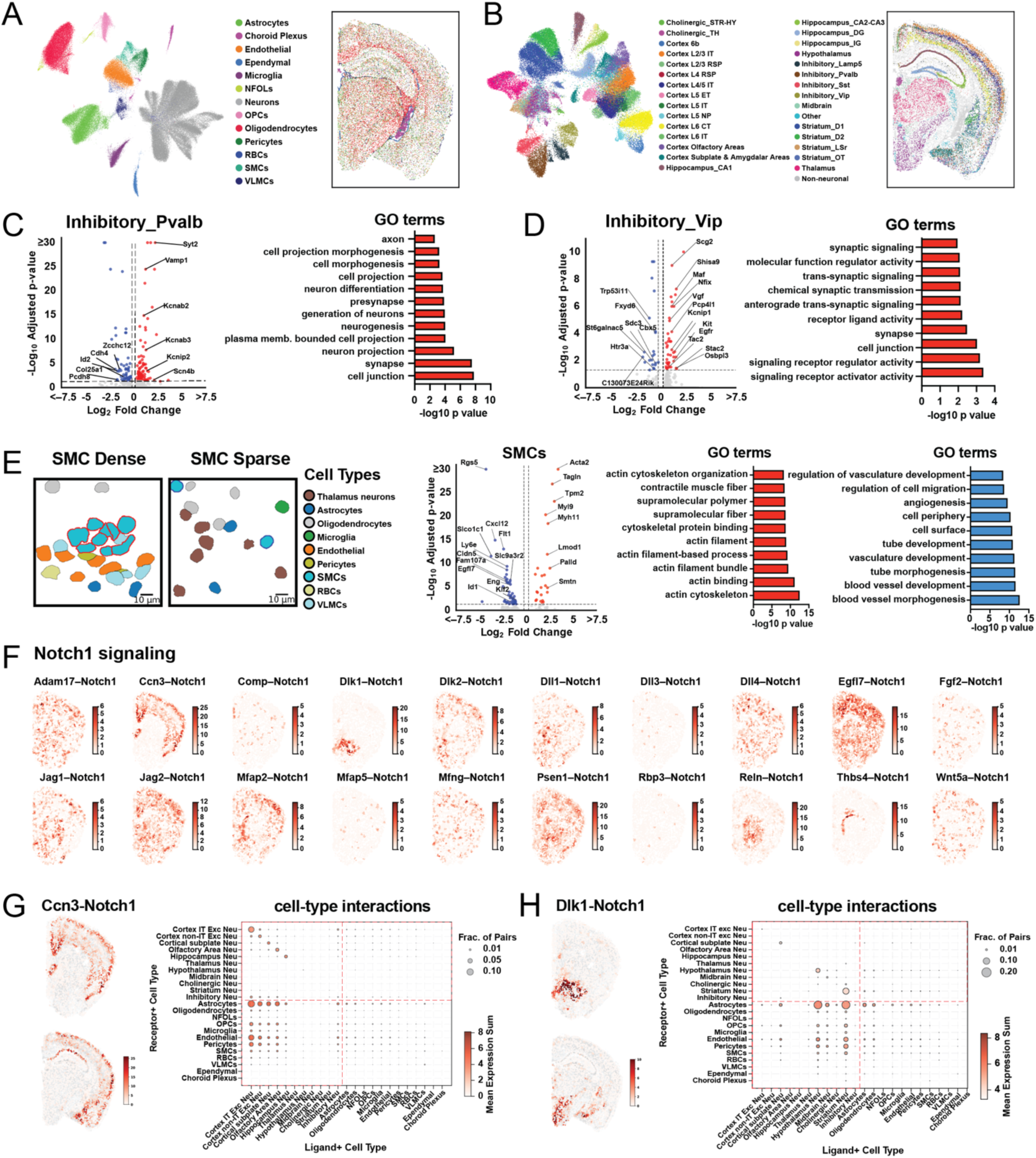
Spatial differential expression (DE) analysis of genes in specific cell types and region- and cell-type-specific ligand-receptor signaling analysis. (A) UMAP of all imaged cells colored by their major cell class identity and spatial map of the major cell classes of a representative posterior section. (B) UMAP of all imaged neurons colored by the subtype identity and spatial map of the neuronal subtypes of the same section as in (A). (C) Spatial DE gene analysis comparing *Pvalb*-positive inhibitory neurons in the isocortex versus all other imaged brain regions. Left: volcano plot showing genes significantly upregulated and downregulated in the isocortex (FDR < 0.05, |log₂FC| > 0.25). Right: selected Gene Ontology (GO) terms enriched in upregulated genes. (D) Same as (C) but for *Vip*-positive inhibitory neurons. (E) DE gene analysis comparing SMCs in densely packed versus sparsely distributed regions, defined by the number of same-type neighbors within a 100 µm radius. Left: Spatial map of a small region showing densely packed SMCs (cyan cells outlined in red) and sparsely distributed SMCs (cyan cells outlined in blue). Middle: Volcano plot showing genes enriched in dense (red) and sparse (blue) SMC cells. Right: GO terms enriched in genes upregulated in densely packed (red) and sparsely distributed (blue) SMCs. (F) Spatial maps of 20 imaged ligand–receptor pairs involving the Notch1 receptor, in a representative anterior brain slice. The spatial maps show, for each receptor-positive cell, the number of ligand-positive cells within a 100 µm radius. (G) Spatial and cell-type distributions of the colocalized Ccn3–Notch1 ligand–receptor pair. Left: Spatial maps of the colocalized Ccn3–Notch1 pair in an anterior slice and a posterior slice. Color scale as (F). Right: Dot plots indicate the sum of ligand and receptor expression (dot color) and the fraction of total interacting cell pairs for this LR pair that belongs to each cell-type pair (dot size). This latter metric evaluates the contribution of each cell-type pair to the observed LR interaction. (H) As in (G), but for the Dlk1–Notch1 pair.

We next investigated how gene expression of the same cell type varied across brain regions using the isocortex versus non-cortical regions as a representative comparison (**Table S4**). Notably, *Pvalb*-positive, *Sst*-positive, *Lamp5*-positive, and *Vip*-positive GABAergic neurons exhibited substantial transcriptional differences between the isocortex and non-cortical regions, containing 183, 109, 78 and 63 significant differentially expressed (DE) genes, respectively. Among non-neuronal populations, mature oligodendrocytes exhibited the greatest transcriptional difference between cortex and non-cortical regions (1,000 DE genes), followed by astrocytes (240 DE genes), microglia (59 DE genes), endothelial cells (43 DE genes), and OPCs (36 DE genes). NFOLs, VLMCs, SMCs, pericytes, and RBCs showed relatively few DE genes (<20 DE genes).

Here, we describe results for two GABAergic neuronal subtypes as examples. In *Pvalb*-positive inhibitory neurons, genes upregulated in the isocortex are involved in cell junction and synaptic transmission, neuronal and axon projection, and neurogenesis and differentiation (**Figure 4C**). Examples genes include potassium and sodium channel subunits (*Kcnab2/3*, *Kcnip2*, *Scn4b*) and synaptic vesicle machinery (*Syt2*, *Vamp1*), which support rapid neurotransmission and suggest enhanced excitability and synaptic activity of *Pvalb* neurons in the isocortex. In *Vip*-positive inhibitory neurons, upregulated genes in the isocortex are involved in synaptic signaling and receptor-ligand activity (**Figure 4D**). Example genes include signaling molecules (e.g. *Osbpl3*, *Kit*, and *Egfr*), genes involved in synaptic modulation (e.g. *Shisa9*, *Kcnip1*, *Pcp4l1*, and *Stac2*), and neuropeptide-related genes (e.g. *Tac2*, *Vgf*, and *Scg2*), suggesting regional variations in synaptic transmission and peptidergic signaling of these neurons.

In addition to capturing transcriptional variations across brain regions, our data also provided insights into how gene expression in individual cell types were affected by their local microenvironment. We illustrate this using SMCs as an example. As a component of the blood vessel wall, SMCs are critical for blood-brain-barrier integrity and can shift phenotypes in response to changing environmental cues.^58^ SMCs exhibit density variations along blood vessels,^59, 60^ forming tightly packed layers in some regions and sparse distributions elsewhere (**Figure 4E**).^61^ Our DE gene analysis showed that densely packed SMCs upregulated gene involved in actin cytoskeleton and contractile function (e.g. *Acta2*, *Myh11*, *Tagln*, *Tpm2*, and *Smtn*) (**Figure 4E**). In contrast, sparsely distributed SMCs showed upregulation of genes associated with vascular remodeling and development, angiogenesis, and signaling (e.g. *Cxcl12*, *Egfl7*, *Id1*, *Fam107a*, and *Klf2*) (**Figure 4E**), suggesting a more adaptive, plastic state suited for vascular restructuring and environmental responsiveness. These results suggest that SMCs adopt context-dependent functional states, ranging from contractile roles in densely packed regions to remodeling and signaling functions in sparse distributed environments.

### Cell-cell communications and ligand-receptor signaling

Spatially resolved transcriptome profiling of single cells also allows for predictions of cell-cell communications based on ligand-receptor (LR) analysis of spatially colocalized cells. We performed spatial proximity/colocalization analysis of ligand- and receptor-expressing cells. Our data include ∼1,500 of the ∼2,000 known LR pairs curated in the CellTalkDB.^62^ For each pair, we calculated an LR colocalization score, defined as the fraction of the total ligand- or receptor-positive cells that have a neighboring cell that is receptor or ligand-positive (**Table S5**).

Top-scoring LR pairs included ligands and receptors involved in diverse developmental and functional pathways. An example top-ranking pair is the platelet-derived growth factor pair (*Pdgfb*-*Pdgfrb*), with the Pdgfb ligand most notably expressed in endothelial cells and the Pdgfrb receptor most notably expressed in pericytes (**Figure S5A**), consistent with the known role of *Pdgfb*-*Pdgfrb* signaling in vascular-cell communication and blood-brain barrier integrity.^63, 64^ Interestingly, we also observe colocalization of *Pdgfrb*-expressing astrocytes with *Pdgfb*-expressing endothelial cells (**Figure S5A**). Since astrocytes are known to play a role in the blood-brain barrier,^59^ our observation suggests that *Pdgfb*-*Pdgfrb* signaling between astrocyte and endothelial cells may also be involved in blood-brain barrier maintenance or regulation.

Another example of top-scoring LR pair is the dynorphin (*Pdyn*) - kappa-opioid receptor (*Oprk1*) pair (**Figure S5B**), which is part of an opioid signaling pathway.^65^ We observed colocalization of this LR pair in the striatum, with *Pdyn* primarily expressed in D1-type MSNs and *Oprk1* expressed in both D1- and D2-type MSNs, consistent with previously described lateral inhibition between D1- and D2-type MSNs via *Oprk1*-mediated suppression of GABA release.^65, 66^

In addition to recovering known cell-cell communications as a validation, our data also provided new insights or hypotheses of cell-type-and region-specific ligand-receptor signaling. To illustrate this, we leveraged the large LR pair coverage of our data to examine families of LR pairs, using the Notch1 signaling pathway as an example. Our data include 20 LR pairs involving the Notch1 receptor (**Figure 4F**; **Figure S5C**). Notably, we observed a highly distinct spatial patterns for different ligand-Notch1 receptor pairs. For example, the *Ccn3*-*Notch1* pair was primarily colocalized in the isocortex, cortical subplate, and olfactory areas, with the *Notch1* receptor most notably expressed in astrocytes and endothelial cells and the *Ccn3* ligand in excitatory neurons (**Figure 4G**). By contrast, the *Dlk1*-*Notch1* pair was colocalized predominantly in the striatum and hypothalamus, with *Notch1* expressed most notably in astrocytes and *Dlk1* in neurons (**Figure 4H**). Although *Ccn3* and *Dlk1* have been implicated in *Notch1* pathway modulation in the brain, little is known about the distinct spatial organizations of astrocyte-neuron *Notch*1 signaling related to these ligands. ^67, 68^ *Ccn3* and *Dlk1* are both non-canonical ligands that can modulate *Notch* signaling, functioning as either activators or inhibitors.^69–71^ The highly distinct spatial distributions of the colocalized *Notch1*-*Ccn3* and *Notch1*-*Dlk1* pairs that we observed thus suggest region-dependent modulation of astrocytic activity and function by neurons via the Notch1 signaling pathway.

### Isoform-resolved single-cell spatial transcriptomics

Whole-transcriptome-scale imaging of single cells provide a powerful framework for analyzing gene regulatory mechanisms and cellular interaction networks in tissues. To further dissect transcriptomic complexity, isoform-level resolution is essential for capturing the functional diversity that arises from alternative splicing, TSS usage, and other transcript variations.^7, 72^ Different isoforms of the same gene could be enriched in distinct cell types or tissue regions and can perform distinct functions. However, isoforms often differ by only short sequence segments at variable sites of the transcripts, hence distinguishing different isoforms remains a major challenge for spatial transcriptomics methods.

Leveraging the ability of RT&T-AMP-MERFISH to detect short transcript sequences, we sought to perform transcriptome-scale single-cell imaging with isoform resolution in the brain, which has a high diversity of splice variants, with alternative splicing playing important roles in brain function and dysfunction in diseases.^72, 73^ In addition to the probe libraries targeting the ∼23,000 genes as described above, we designed an additional probe library targeting ∼10,000 annotated isoforms for ∼4,000 of the ∼23,000 genes, comprising splice variants as well as isoforms differing in TSS or other transcript features. In order to image these ∼10,000 isoforms together with the ∼23,000 genes (∼33,000 distinct transcript sequences in total), we added an additional MERFISH run designated to detecting the isoforms, also using a 90-bit Hamming-distance 4 and Hamming-weight 4 code. In total, three back-to-back MERFISH runs were performed on each sample, using 90 rounds of three-color imaging, to detect the ∼33,000 transcript sequences. As for the 23,000 genes, we also used 6 encoding probes per isoform, targeting the unique sequences in each isoform. We imaged three coronal brain slices at a posterior position (∼130,000 cells in total). We observed excellent reproducibility of the RT&T-AMP-MERFISH data between replicates and good correlation with bulk RNA sequencing data (**Figure 5A-C**).

**Figure 5.**
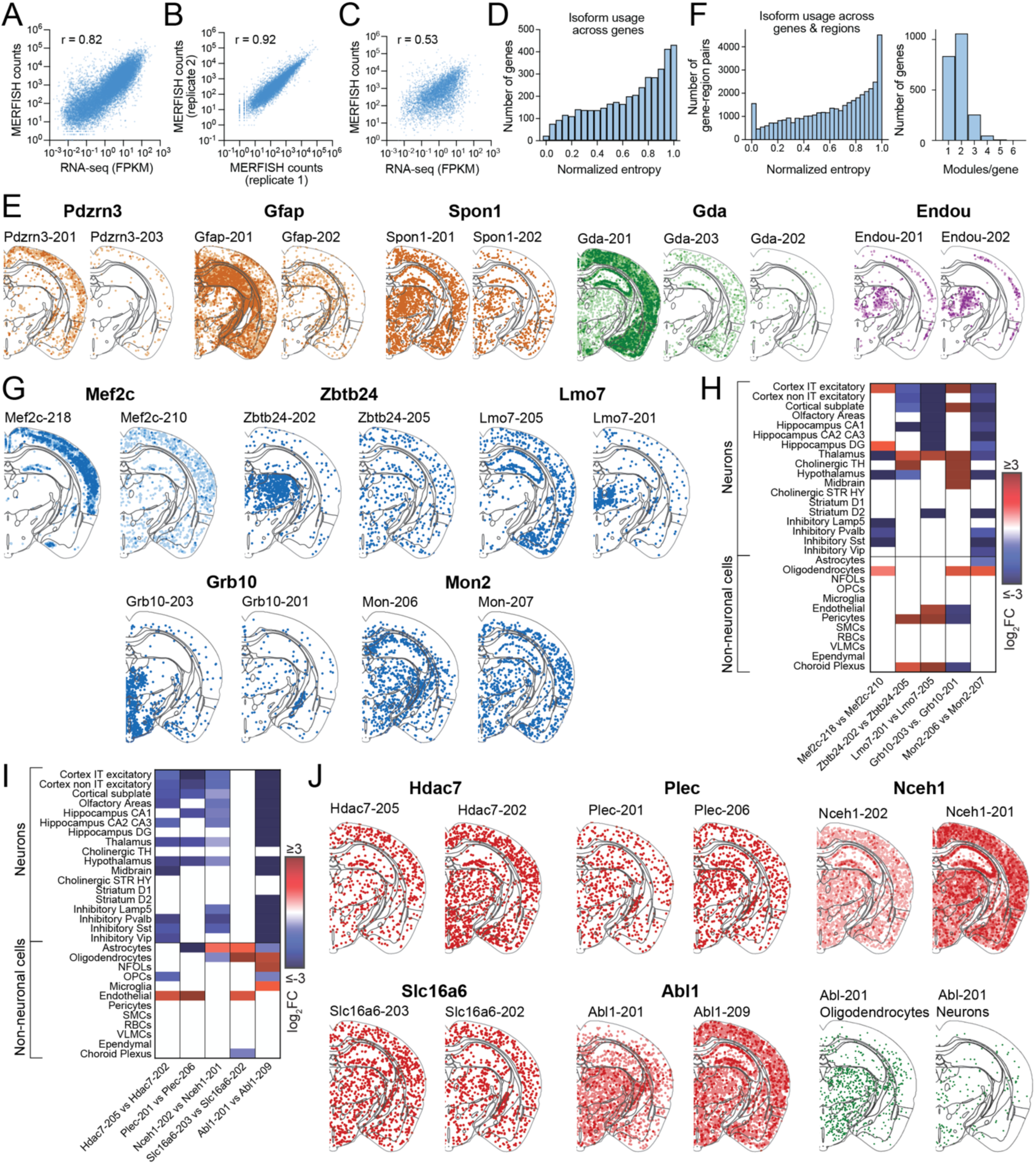
Isoform-resolved, single-cell spatial transcriptomics with RT&T-AMP-MERFISH reveals diverse region-and cell-type-specific isoform usage patterns in the mouse brain. (A) Correlation between the expression levels determined by RT&T-AMP-MERFISH and bulk RNA sequencing for the ∼23,000 genes (imaged together with the ∼10,000 isoform-defining transcripts). (B) Reproducibility between replicates for the expression levels determined by RT&T-AMP-MERFISH for the ∼10,000 isoforms. (C) Correlation between the expression levels determined by RT&T-AMP-MERFISH and bulk RNA sequencing for the ∼10,000 isoforms. (D) Distribution of the normalized Shannon entropy for isoform expression across all measured genes over the entire measured samples. For each gene, the Shannon entropy quantify the overall isoform usage bias, with values range from 0 (expression dominated by a single isoform) to 1 (equal usage of all isoforms). (E) Representative examples of overall isoform usage patterns ranging from highly biased expression of one isoform to balanced expression of all isoforms. *Pdzrn3*, *Gfap*, and *Spon1* showed strong bias toward a single isoform; *Gda* showed graded expression across multiple isoforms; *Endou* showed balanced usage between two isoforms. The grey lines in (E), (G) and (J) demarcate brain regions defined in the Allen CCF. (F) Left: Normalized Shannon entropy as in (D) but with analysis performed on individual brain regions defined in Allen CCF v3. Results across all regions and all genes are shown in one histogram. Right: Isoform-level spatial Hotspot analysis showing the number of spatial modules assigned to each gene based on the spatial patterns of its isoforms. Genes with no isoform assigned to any spatial module (i.e. exhibiting no specific spatial patterns) are not included in the plot. (G) Spatial maps showing the distribution of two isoforms for each of five example genes (*Mef2c*, *Zbtb24*, *Lmo7*, *Grb10*, and *Mon2*) that show region- and cell-type-specific isoform usage. (H) Heatmap corresponding to the genes in (G), showing log₂ fold change between the fractions of cells expressing the two isoforms for indicated cell types. (I) Same as (H), but for five different genes *Hdac7*, *Plec*, *Nceh1*, *Slc16a6*, and *Abl1*, that show cell-type-specific isoform usage, albeit with largely uniform spatial distribution. (J) Spatial maps corresponding to the genes in (I). Bottom right shown in green: Spatial maps for the *Abl*-201 isoform in the oligodendrocyte-lineage cells (left) and in neurons (right).

We first evaluated isoform usage patterns for each gene in terms of the overall expression level in a cell-type or brain-region independent manner. To this end, we determined the fractional abundance of each isoform relative to the gene’s total expression across the whole samples and computed the Shannon entropy based on these relative expression levels across different isoforms (**Methods**). To enable comparisons across genes with differing isoform numbers, we normalized each gene’s entropy by the theoretical maximum entropy achievable for that number of isoforms (**Methods**). This normalized entropy metric ranges from 0 (complete dominance by one isoform) to 1 (equal usage of all isoforms), providing a continuous measure of isoform usage across genes.

Across all genes analyzed, we observed a broad spectrum of isoform usage patterns (**Figure 5D**). Some genes showed highly uneven expression dominated by a single isoform, which was reproducible across replicates. Three such examples are shown in **Figure 5E**, including *Pdzrn3*, (encoding a PDZ domain-containing RING finger protein), *Gfap* (encoding the glial fibrillary acidic protein), and *Spon1* (encoding an extra-cellular matrix protein), with *Pdzrn3*-201, *Gfap*-201, and *Spon1*-201 being the dominantly expressed isoforms. Consistent with and validating our results, *Spon1-201* and *Gfap*-201 have also been shown previously to be the dominant isoforms of their respective genes in the cortex.^74, 75^ In addition to such patterns with single-isoform dominance, we also observed more graded patterns where multiple isoforms of the same gene showed increasingly higher expression (e.g. *Gda*, which encodes a guanine deaminase) and more balanced patterns where different isoforms of the same gene showed nearly equal expression (e.g. *Endou*, an endonuclease gene) (**Figure 5E**). Genes in this latter group often exhibited similar spatial distributions (like *Endou*), which agree with the gene-level spatial expression pattern observed in Allen Brain Atlas in situ hybridization data,^76^ again validating our RT&T-MAP-MERFISH results.

### Region- and cell-type-specific isoform usage

Our isoform-resolved spatial transcriptomics data with single-cell resolution further allowed examination of isoform diversity across both brain regions and cell types. To assess spatial variations in isoform usage, we applied two quantitative analyses as an initial screen. In the first analysis, we computed the Shannon entropy for each gene, as described above, but now separately for each brain region assigned based on the Allen CCF (**Figure 5F, left panel**). This entropy distribution across gene-region pairs again exhibited a broad distribution, but with a bias towards “0” entropy as compared to the region-agnostic distribution (compare **Figure 5F** with **Figure 5D**), indicating that some genes have strong isoform selection in individual brain regions. In the second analysis, we performed the spatial Hotspot analysis at the isoform level, assigning each isoform to a spatial module based on its spatial expression pattern. A gene with more than one spatial module assignments indicates that at least some of the different isoforms were assigned to different modules (**Figure 5F, right panel**). Notably, many genes had their different isoforms (or different subsets of isoforms) assigned to different spatial modules, suggesting different spatial distributions for these isoforms. From these analyses, we screened for genes with entropy values substantially lower than 1 and/or module/gene numbers greater than 1 for further examination and describe several interesting examples below.

Our analyses revealed a high diversity of isoform expression patterns across genes, in terms of both brain-region and cell-type specificity. **Figures 5G and 5H** show five of the many genes that exhibited brain-region- and cell-type-specific isoform expression patterns, which were reproducible across replicates. One notable example is *Mef2c*, which expressed a protein-coding isoform and a non-coding isoform with strikingly distinct spatial distributions and cell-type specificity. The protein-coding isoform (*Mef2c*-218) encodes a calcium-responsive transcription factor that activates genes involved in neuronal development.^77^ Dysregulation of this isoform has been linked to neurodevelopmental disorders, including autism and schizophrenia.^78^ In our data, this coding isoform of *Mef2c* was enriched in excitatory neurons in the cortex and dentate gyrus (**Figure 5G, H**), both regions associated with cognitive function, consistent with previous knowledge of where this transcription factor is expressed and performs its function.^77, 79^ In contrast, we observed that the non-coding isoform *Mef2c*-210 was broadly expressed in many brain regions (e.g. cortex, thalamus, hypothalamus, etc.) and, in the cortex and dentate gyrus, this isoform was specifically enriched in inhibitory neurons (**Figure 5G, H**). Although the function of the non-coding isoform for *Mef2c* is unknown, for some other genes, non-coding isoforms can regulate the expression of their protein-coding counterparts, typically through repression mechanisms such as competition for promoter or splicing machinery.^80–83^ The expression patterns that we observed for the two *Mef2c* isoforms thus suggests an interesting regulatory model in which the non-coding isoform modulates Mef2c expression, suppressing the expression of the protein-coding isoform in inhibitory neurons in the cortex and dentate gyrus as well as in neurons in other brain regions, leading to specific expression of the protein-coding isoform, and hence the Mef2c protein, in the excitatory neurons of the cortex and dentate gyrus for its function.

Other examples also showed striking spatial and cell-type-dependent isoform expression pattern (**Figure 5G, H**). For example, *Zbtb24*, a transcription factor gene involved in epigenetic regulation and DNA methylation,^84^ had one isoform (*Zbtb24*-202) expressed nearly exclusively in the thalamus neurons, while the second isoform (*Zbtb24*-205) was broadly expressed across many different brain regions. *Lmo7*, which encodes a multifunctional scaffold protein that connects the actin cytoskeleton to cell-cell adhesion complexes and transmits mechanical signals to the nucleus for regulating gene expression,^85^ expressed seven isoforms with diverse spatial patterns (shown later in **Figure S6C**). Two of the isoforms are shown in **Figure 5G, H**: one isoform (*Lmo7*-201) was primarily expressed in the medial part of the thalamus, while the other (*Lmo7*-205) was primarily expressed in the cortex, cortical subplate, olfactory areas, and hippocampus. Grb10, which is involved in growth factor signaling,^86^ had one isoform (*Grb10*-203) strongly enriched in the medial parts of thalamus and hypothalamus and another isoform (*Grb10*-201) strongly enriched in the cells of the choroid plexus and endothelial cells. Mon2, a gene involved in vesicle transport,^87^ had one isoform (*Mon2*-206) primarily expressed in oligodendrocytes, particularly enriched in the fiber tract, while the second isoform (*Mon2*-207) was broadly expressed in neuronal cell types across many brain regions.

It is also worth noting that some genes, albeit not exhibiting highly distinct spatial patterns for different isoforms, nonetheless showed strong cell-type specificity in isoform expression, in particular for non-neuronal cells (**Figure 5I, J**). For example, Hdac7, a Histone Deacetylase involved in transcriptional regulation,^88^ had one isoform (*Hdac7*-205) enriched in endothelial cells and the other isoform (*Hdac7*-202) enriched in neurons. Interestingly, Hdac7 isoforms have been implicated in various cancers,^89^ and the specific isoform expression patterns may have implications for different cancers. Likewise*, Plec*, a gene encoding Plectin and contributing to cytoskeletal organization and cell-cell junction stability,^90^, also had one isoform (*Plec*-201) preferentially expressed in endothelial cells and the other isoform (*Plec*-206) enriched in some neuronal populations. *Nceh1*, a gene encoding a cholesterol ester hydrolase enzyme,^91^ had one isoform (*Nceh1*-202) enriched in astrocytes and the other isoform (*Nceh1*-201) enriched in neurons. Slc16a6, a member of the monocarboxylate transporter family,^92^ had one isoform (*Slc16a6*-203) enriched in oligodendrocytes, astrocytes, and endothelial cells, while another isoform (*Slc16a6*-206) was enriched in cells of the choroid plexus. Abl1, a tyrosine kinase involved in cell division, growth, and migration,^93^ had a protein-coding isoform (*Abl1*-201) expressed in oligodendrocytes, while a non-coding isoform (*Abl1*-209) was enriched primarily in neuronal cell types. Interestingly, the protein-coding isoform *Abl1*-201 further showed brain-region specificity, with enrichment in oligodendrocytes in the fiber tract, thalamus, and hypothalamus.

Overall, our data showed a remarkably rich diversity of isoform usage patterns of genes across cell types and anatomical regions in the brain, demonstrating the power of RT&T-AMP-MERFISH for revealing alternative splicing and other transcript variations with high spatial and cell-type resolution.

### Brain structures enriched for specific isoform usage across many genes

Notably, our data revealed that some brain regions and structures exhibited a particularly high frequency of isoform switching across genes, i.e. a particularly large number of genes showed specific isoform usage in these structures. Two of the most notable examples were the choroid plexus (CP) and the hippocampus. Among the ∼1,400 genes that we observed to have isoforms assigned to more than one spatial gene module, indicating spatially dependent isoform usage, ∼100 genes had one of their modules corresponding to the CP and ∼800 genes had a subset of their modules corresponding to CA1, CA2/3, or DG of the hippocampus.

### Choroid plexus (CP)

The CP is a network of blood vessels residing in the brain’s ventricles, and CP epithelial cells form a specialized barrier between the blood and cerebrospinal fluid (CSF), producing and regulating CSF. Notably, our data showed that numerous genes had highly specific isoform usage in CP cells of the adult mouse brain. In **Figure 6A, B** and **Figure S6A, B**, we show some examples of the genes exhibiting isoform specificity in CP cells, including *Csnk1e*, *Crhr2*, *Slc31a1*, *Sept9*, *Dlg3*, *Ace*, *Ccbe1*, *Coq8b*, *Enpp5*, *Lamp2*, *Ogdh*, *Ppp1r1b*, *Tbc1d9*, and *1500011B03Rik*. These genes span multiple functional classes, suggesting that CP cells may tailor protein function through isoform usage to support their specialized roles in blood-brain barrier function, circadian clock regulation, and signaling.

**Figure 6.**
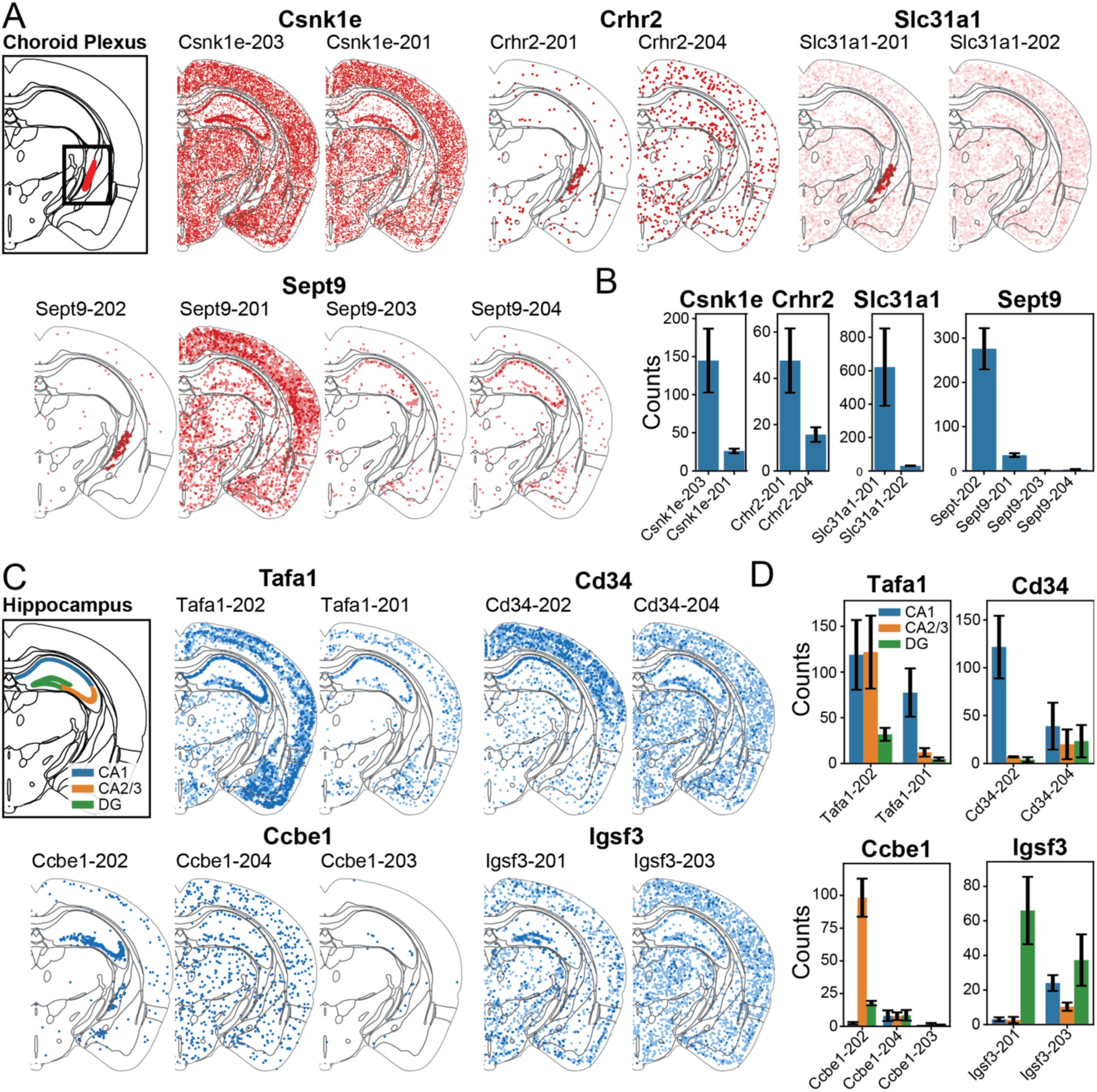
Two brain structures that exhibit specific isoform usage for numerous genes. (A) Allen CCF showing the choroid plexus (CP) and spatial maps of example genes (*Csnk1e*, *Crhr2*, *Slc31a1*, and *Sept9*) exhibiting strong specificity for isoform usage in the CP. More examples are shown in **Figure S6A, B**. The grey lines in demarcate brain regions defined in the Allen CCF. (B) Expression levels (in mean ± SEM of counts, across replicates) in CP cells for the different isoforms of the genes shown in (A). (C) Allen CCF showing the CA1, CA2/3, and DG in hippocampus and spatial maps of example genes (*Tafa1*, *Ccbe1*, *Cd3*4, and *Igsf3*) exhibiting strong specificity for isoform usage in the hippocampus or subregions of the hippocampus. More examples are shown in **Figure S6C, D**. (D) Expression levels (in mean ± SEM of counts, across replicates) in CA1, CA2/3, and DG cells for the different isoforms of genes shown in (C).

We briefly describe a few examples here. One notable example is *Csnk1e*, which encodes a core circadian clock kinase (CKIe).^94^ We imaged two protein-coding isoforms of *Csnk1e*. While both isoforms were broadly expressed across many brain regions, only one of them (*Csnk1e*-203) was substantially expressed in CP cells (**Figure 6A, B**). Interestingly, the CP has a peripheral circadian clock that differs from the master circadian clock in the suprachiasmatic nucleus (SCN) and functions to tune CSF composition, supporting time-of-day-dependent processes such as waste clearance, nutrient delivery, and neuroendocrine signaling.^95–99^ CKIe proteins encoded by the two *Csnk1e* isoforms have identical kinase domains, but differ in their C-terminal domain important for substrate specificity and kinase activity.^100^ Our observations thus suggest that this structural variation and its associated substrate specificity could be important for regulating the CP-controlled circadian clock.

Another example is *Crhr2*, which encodes a GPCR involved in neuroendocrine and stress-related signaling.^101, 102^ We observed that one of the two imaged *Crhr2* isoforms (*Crhr2*-204) was expressed broadly across brain regions and cell types, whereas the other (*Crhr2*-201) was nearly exclusively expressed in CP cells (**Figure 6A**) at a level much stronger than that of *Crhr2*-204 (**Figure 6B**). The two isoforms differ in their 3′ ends of the coding sequence. In GPCRs, the C-terminal region is generally important for trafficking, internalization, and signaling interactions, and thus changes in this region can alter receptor function. The enrichment of the *Crhr2*-201 isoform in CP cells suggests the possibility of a CP-specific role of this GPCR facilitated by its C-terminal region.

*Slc31a1*, which encodes a copper transporter, ^103^ also exhibited high specificity of isoform usage. *Slc31a1*-202 was broadly expressed across many brain regions, whereas *Slc31a1-201* was strongly enriched in CP cells, expressing at a level >10-fold higher than *Slc31a1*-202 (**Figure 6A, B**). The CP-enriched isoform is transcribed from a different promoter and contains an additional exon that substantially alters the protein structure. The predominance of this *Slc31a1* isoform in the CP suggest a specific role of this isoform in copper transport, which could be of medical importance as the CP plays a central role in brain copper metabolism^104^ and deficiencies in CP-mediated copper transport are linked to neurological disorders.^105^

Cytoskeleton-related proteins likewise displayed region- and cell-type-specific isoform expression patterns, exemplified by *Sept9*, which encodes a septin protein. Of the four isoforms of *Sept9* that we imaged, *Sept9*-201 was broadly expressed across brain regions, *Sept9*-203 and *Sept9*-204 were enriched in hippocampal neurons, while *Sept9*-202 was highly enriched in the CP (**Figure 6A, B**). Septins are cytoskeletal GTP-binding proteins that play important roles in cell division, membrane remodeling, and vesicle trafficking.^106^ Isoforms of *Sept9* have been identified and studied in several tissue types,^107, 108^ but region- and cell type-specific roles of *Sept9* isoforms in the brain have yet to be characterized. Our findings offer an entry point to investigate the specific role of this protein in CP function.

It is interesting to note that alternative splicing and isoform diversity are particularly prominent and dynamically regulated during development, with many isoforms being down- or up-regulated as the organ matures.^109–111^ In the developmental brain, CP epithelial cells are highly enriched for differential isoform expression.^13^ Given our results that numerous genes exhibit highly specific isoform usage in CP cells of the adult brain, an interesting question for future investigation would be how isoform usage in the CP is regulated and evolves as this important brain structure matures.

### Hippocampus

Numerous genes showed specific isoform usage in hippocampal neurons. Some of the examples are shown in **Figure 6C**, **D** and **Figure S6C, D**, including *Mef2c* (as described earlier), *Tafa1*, Ccbe1, *Cd34*, *Igsf3*, *Actr3b*, *Abhd2*, *Adarb2*, *Epha6*, *Lmo3*, *Pantr1*, *Prkag2*, *Smarcc2* and *Lmo7*. Among these, some genes showed enrichment of specific isoforms across all four subregions, whereas many genes showed specific isoform enrichment only in some subregions of the hippocampus (e.g. *Tafa1*, Cd34, and *Lmo3* in CA1; *Ccbe1* in CA2/3; *Epha6* in CA1-CA3; *Igsf3* and *Mef2c* in DG).

For example, the two imaged *Tafa1* isoforms showed distinct expression patterns across hippocampal subregions. *Tafa1*-202 was enriched in CA1-CA3 as compared to DG, whereas Tafa1-201 was enriched in CA1 (**Figure 6C, D**). Both isoforms were also expressed in the isocortex, cortical subplate, and olfactory areas (**Figure 6C, D**). Structural comparison showed that the protein-coding sequence of the two isoforms was identical, but their 5’ UTR, TSS, and 3′ UTR differ. These features point to differences in transcriptional and post-transcriptional regulation as the likely basis for their distinct expression profiles. Given that *Tafa1* encodes a brain-enriched, chemokine-like protein implicated in dendritic spine maturation and synaptic function,^112^ isoform-specific regulation in the CA1-CA3 and DG neurons may influence region-specific synaptic activity and plasticity in the hippocampus.

Many of the hippocampus-specific examples involved different usage of protein-coding and non-coding isoforms. As described above, the *Mef2c* protein-coding isoform was specifically enriched in DG excitatory neurons (as well as cortical excitatory neurons), whereas its non-coding isoform was expressed in other neuronal cell types (**Figure 5G, H**; **Figure S6C**, **D**). Likewise, *Ccbe1* (encoding an extracellular matrix protein), *Cd34* (encoding a cell surface glycoprotein), and *Igsf3* (encoding a cell adhesion molecule) each had a protein-coding isoform enriched in subregions of the hippocampus: *Cd34*-202 in CA1 (also in cortex), *Ccbe1*-202 in CA2/3, and *Igsf3*-201 in DG (**Figure 6C**, **D**). In contrast, their non-coding isoforms, often containing retained intronic sequences, were expressed across many brain regions. Intron retention can regulate gene expression through transcript stability, nuclear retention, and translation efficiency.^80–83, 113–115^ Increased intron retention in the hippocampus has been reported in aging and Alzheimer’s disease.^113, 116–118^ Our findings thus indicate that isoform-specific regulation, including the production of non-coding variants, contributes to region-specific molecular specialization within the hippocampus, suggesting a possibility that dysregulation in isoform expression could cause hippocampus-related neurological disease.

Together, our data illustrate that isoform-resolved spatial transcriptomic profiling of single cells can reveal novel molecular specializations underlying distinct brain regions and cell types. While the functional importance of our observed region- and/or cell-type-specific isoform usage patterns awaits future investigations, our data provide a resource and entry point for such functional investigation by pointing to the specific isoforms to perturb genetically, for example via the RNA-targeting CRISPR-Cas13 system,^119^ and the specific cell types and brain regions in which to perform these genetic perturbations.

## DISCUSSION

In this work, we developed a spatial transcriptomics method based on a novel *in situ* RNA amplification approach (RT&T-AMP) and its integration with MERFISH. Our method enabled isoform-resolved and whole-transcriptome-scale gene expression profiling of single cells with high spatial resolution in intact tissues. This amplification scheme allows detection of essentially all genes, including short genes that are difficult to measure by traditional multiplexed FISH. Here, we measured ∼23,000 genes. The remaining few thousand expressed genes were not imaged because of their very high expression levels, but could also be included by adding additional MERFISH runs to the same samples. In parallel to our work, recent developments reported in bioRxiv have demonstrated imaging of ∼20,000 genes by integrating branched DNA amplification or RCA with multiplex FISH,^120, 121^ but transcriptome imaging with isoform resolution has yet to be demonstrated. Here, we demonstrated isoform-resolved spatial transcriptomics of single cells using RT&T-AMP-MERFISH. Leveraging this ability, we imaged ∼10,000 isoform-defining transcripts together with the 23,000 genes (measuring ∼33,000 distinct RNA sequences total), representing the largest transcript set measured by imaging to date. Our smFISH data further suggest the feasibility of using as few as a single probe for imaging, which should allow most isoforms to be resolved. Moreover, compared to unamplified MERFISH, our method reduced probe cost by an order of magnitude, which should further facilitate the broad adoption of whole-transcriptome-scale imaging.

By capturing all or nearly all expressed genes, RT&T-AMP-MERFISH allows essentially unbiased, gene expression profiling while maintaining single-cell and subcellular resolution. This capability allows comprehensive cell-type and cell-state classification, systematic differential gene expression analysis across cell types/states and tissue regions, and large-scale predictions of cell-cell communication and ligand-receptor signaling. In this work, we performed spatial differential gene expression analyses in mouse brain tissue, identifying spatially regulated gene programs, including those modulated by the local microenvironment. We also identified gene modules associated with anatomical structures in the brain, generating hypotheses of functional connections between genes. Taking advantage of the large transcriptomic coverage and high spatial resolution, we also conducted interaction analyses across nearly all expressed ligand-receptor pairs, providing insights into region- and cell-type-specific cell-cell communications and ligand-receptor signaling.

Notably, our isoform-resolved, single-cell transcriptome imaging enables systematic analysis of isoforms arising from alternative splicing and other transcript variations. Leveraging this capability, we identified region- and cell-type-specific isoform usage for numerous genes in the adult mouse brain, as well as brain structures particularly enriched for specific isoform usage across a large number of genes. Our data not only revealed a remarkable level of spatial diversity and cell-type specificity in alternative splicing and other transcript variations, but also provide a rich resource for future functional interrogations of these isoform usage patterns by pointing to the specific isoforms to perturb in specific cell types and brain regions. Overall, our data illustrate how isoform-resolved, whole-transcriptome-scale imaging unlocks new discoveries that are challenging with previous methods.

Beyond messenger RNAs and long non-coding RNAs (lncRNAs), many other RNA species such as miRNAs, siRNAs, piRNAs, snRNAs, and snoRNAs, also perform critical cellular functions, including transcriptional, post-transcriptional, and translational gene regulation.^122, 123^ RT&T-AMP could potentially also make these small RNAs accessible by transcriptome imaging, for example by using *in vitro* polyadenylation to add poly-A tails to these RNAs,^124^ which could then enable poly-T primed reverse transcription followed by transcription for *in situ* amplification, as in RT&T-AMP. We thus anticipate that RT&T-AMP-facilitated transcriptome imaging will not only allow us to image all genes, including their distinct isoforms, but also other functionally important cellular RNAs, providing transcriptome-wide expression profiling of single cells with high spatial resolution, opening new pathways of discoveries in cell and tissue biology.

### Limitations of the study

In this work, we measured ∼23,000 genes and ∼10,000 isoforms. There are many additional annotated isoforms that we did not measure. Our method should allow us to include many of these isoforms in future studies. In addition, we imaged only several sections of the brain, which does not provide a full sampling of brain cell types and anatomic structures. Future isoform-resolved, whole-transcriptome imaging of the entire brain will provide a more complete picture of the molecule and cellular architecture of the brain. Finally, while our approach identified spatial patterns and cell-type specificity of gene and isoform expression, future functional studies will be needed to investigate the biological roles of the spatial and cell-type dependence in the gene and isoform expression patterns observed here.

## ACKNOWLEDGEMENTS

We thank members of the Zhuang lab for helpful discussions. This work is in part supported by the Howard Hughes Medical Institute (HHMI). L.C. is supported in part by a Brightfocus Foundation Postdoctoral Fellowship. Acknowledgement is made to the donors of Alzheimer’s Disease Research, a program of Brightfocus Foundation. X.P. is supported in part by the Jane Coffin Childs - HHMI Fellowship. X.Z. is a HHMI investigator.

## AUTHOR CONTRIBUTIONS

L.C. and X.Z. conceived of the study. L.C., A.H., T.B. and X.Z designed the experiments. T.B. designed the gene and isoform panels and probe libraries. L.C., A.H., and P.C. performed the experiments. L.C., A.H., T.B., and X.P. performed data analysis. L.C., A.H., T.B., P.C., X.P., and X.Z. interpreted data. L.C. and X.Z. wrote the manuscript with input from A.H., T.B., P.C. and X.P. X.Z. supervised the study.

## METHODS

### Brain tissue sample preparation

Brains from male C57BL/6J mice (Jackson Laboratory, JAX #000664) at 14 weeks of age were used in this study. Mouse brains were directly acquired from the Jackson Laboratory and frozen in optimal cutting temperature compound (O.C.T.) and stored at -80 °C until sectioning.

Prior to sectioning, brains were taken from storage at -80 °C and warmed to -20 °C in a cryostat (Leica) for 30 minutes. 10-μm coronal sections were collected and placed on a 40-mm-diameter coverslip (Bioptechs). For RT&T-AMP MERFISH measurements of of ∼23,000 genes, two anterior coronal sections and two posterior coronal sections were imaged. For RT&T-AMP MERFISH measurements of ∼23,000 genes and ∼10,000 isoforms, three posterior coronal sections were imaged.

Tissue sections were fixed with a glyoxal-based fixative, prepared fresh by combining the following components: 850 μL nuclease-free water, 50 μL 100% ethanol, 77.5 μL 40% glyoxal (Sigma 50649), and 7.5 μL glacial acetic acid (Sigma A6283). The pH was adjusted to 4-5 using approximately 6μL of 1 N NaOH. Finally, 10 μL of RNasin (Promega N2611) was added to the solution. The fixative was pre-chilled on ice and then applied to the samples for 15 minutes. The samples were then washed 3 times with 1xPBS and processed for RNA amplification by reverse transcription and transcription (RT&T-AMP) immediately. Tissue samples were washed with nuclease-free water followed by 2xSSC. Next, 100 μL of a poly-T primer (GATGATGGAGGGAATAAGTTTT+TT+TT+TT+TT+TT+TT+TT+TT+TT+TT+TT+TTTT, IDT) solution was added. The solution contained 1 μM of primer in 5xSSC, supplemented with 1:100 RNasin and used in a Silicone Imaging Chamber (EMS, 70327-05). Samples were incubated overnight at 37 °C.

Following overnight incubation, the tissue was washed 3 times with 1xPBS and incubated with 100 μL of reverse transcription (RT) and template switching reaction solution. A 200 μL reaction solution was prepared as follows: 114 μL nuclease-free water, 15 μL dNTP (10mM stock, ThermoFisher R0192), 5 μL template switching oligo (TSO sequence: /5Biosg/TAATACGACTCACTATAGGGAAATA rGrG+G, 100uM stock, IDT), 50 μL RT buffer (5x stock, ThermoFisher EP0753), 3 μL Rnase inhibitor (40 U/μL stock, NEB M0314L), 3 μL RNasin, 10 μL Maxima H minus (200 U/μL stock ThermoFisher EP0753). All components were first added to the reaction solution except for the RT buffer and the enzyme, and the reaction solution was mixed thoroughly. Then the RT buffer was added, mixed thoroughly, and finally the RT enzyme was added and mixed gently. The TSO was reconstituted in TE pH8, aliquoted for single use, and stored at -80 °C. The reaction was incubated in the same Silicone Imaging Chamber overnight at 37 °C, for a maximum of 18 hours. The silicone Chamber was washed 2 times with water first. After overnight incubation, the tissue samples were washed 3 times with 1xPBS, and then the T7 transcription reaction was initiated. The T7 reaction components were as follows: 64 μL nuclease-free water, 3 μL Rnase inhibitor (40 U/μL NEB), 3 μL Rnasin, 100 μL NTP solution (NEB E2050S), 10 μL DTT (NEB E2050S) and 20 μL T7 polymerase (NEB E2050S). All components except for the enzyme were first mixed, and then the enzyme (T7 polymerase) was added last to the reaction solution and mixed gently. 100 μL of reaction solution was carefully placed on the tissue sample and the petri dish was sealed with parafilm. The tissue was incubated for three hours at 37 °C. After the transcription reaction, 100 μL of 4% PFA (ThermoFisher 28906) was placed on the sample carefully, without removing the T7 reaction solution. The samples were incubated for 15 minutes and then washed 3 times with 1xPBS. The samples were then either hybridized with encoding probes or stored in 1xPBS supplemented with 1:1000 RNasin for later hybridization.

For encoding probe hybridization, tissue samples were first incubated with 30% formamide for 5 minutes. Then, 50 μL of hybridization solution for smFISH or 30μL of hybridization solution for whole-transcriptome-scale MERFISH was added to a petri dish covered in parafilm. The tissue sample was then carefully inverted and placed on top of the solution and incubated for 48 hours at 37 °C. The encoding probe hybridization buffer was prepared as follows: 300 μL 100% formamide, 500 μL 20% w/v dextran sulfate (Millipore S4030), 100 μL 20x SSC, 100 μL nuclease-free water, 10 μL yeast tRNA (10mg/mL stock ThermoFisher AM7119), 1 ul Rnasin, 1 uM of (/5Acryd/GATGATGGAGGGAATAAGTTTT+TT+TT+TT+TT+TT+TT+TT+TT+TT+TT+TT+TTTT, IDT), and encoding probes at concentrations described below. smFISH probes were hybridized at 4 μM per probe. The 30μL whole-transcriptome-scale MERFISH hybridization solution included two probe libraries. The first library (codebook 1, ∼20,000 genes) was hybridized at 0.5 nM per probe. The second library (codebook 2, ∼3,000 genes) was hybridized at 1 nM per probe. The third library (codebook 3, ∼10,000 transcripts) was hybridized at 1 nM per probe. Following incubation with the hybridization solution, the sample was washed 2x with 30% formamide at 47 °C, 30 minutes per wash. Then the sample was then washed 3 times with 2xSSC, and then imaged or stored at 4 °C in 2xSSC supplemented with 1:1000 Rnasin before imaging. The two whole transcriptome codebooks were imaged in two back-to-back experiments on the same samples, and the whole transcriptome and isoform codebooks were imaged in three back-to-back experiments on the same samples.

### RT&T-AMP-MERFISH gene, isoform, and probe library design

Genes included in the whole-transcriptome-scale MERFISH library were selected from the Ensembl transcriptome, release 102. From the ∼52,000 unique gene symbols, we first limited our scope to those genes and transcripts with gene or transcript biotype of ‘protein_coding’ or ‘lincRNA’. We next removed histone and mitochondrial genes. We then compared our list of genes with that reported in the Allen Institute’s whole-brain snRNA-seq data ^40^ and proceeded with the intersection of the two lists. This resulted in ∼27,000 unique gene symbols. This list missed 16 genes that were included in our previous whole-mouse-brain MERFISH study,^41^ which we added back.

We next removed genes with very high expression in the mouse brain. First, we removed all genes with expression greater than or equal to that of *Actb* (encoding actin) as reported in bulk RNA sequencing results^41^. Secondly, using the Allen snRNA-seq results,^40^ we determined, for each gene, both its ‘subclass-max’ (maximum expression value across all reported subclasses) and ‘cluster-max’ (maximum expression value across all reported clusters). We then removed all genes with a subclass-max greater than that of *Gnrh1* and all genes with a cluster-max greater than that of *Mef2c*. In each case, the threshold gene selected (*Gnrh1* or *Mef2c*) was the gene of the highest expression level among our previously reported whole-brain MERFISH panel.^41^ In total, the filtering steps to filter out highly abundant genes removed only ∼90 genes.

The remaining ∼27,000 we genes were then divided into two groups of ∼22,000 ‘Tier 1’ genes with relatively low expression level and ∼5,000 ‘Tier 2’ genes with relatively high expression level, where the latter tier was defined as those genes with a subclass-max or cluster-max greater than one half of the value of *Sst* for the corresponding *Sst* positive GABAergic neurons. From Tier 2, we further selected the first 3000 genes when ranked from lowest to highest by their cluster-max expression and included additional genes that were on our previous MERFISH gene panels for various brain regions but were not included in these 3000 genes or Tier 1 genes.

These two tiers of genes were imaged with two back-to-back MERFISH runs with two MERFISH codebook, Codebook 1 and Codebook 2. Each codebook contains a 90-bit Hamming Distance 4 and Hamming Weight 4 code, to be imaged with two orthogonal sets of readout probes. The MERFISH encoding probe design for each tier of genes were performed using a previously reported computational pipeline (Github: https://github.com/xingjiepan/MERFISH_probe_design).^41^ After the encoding probe design, a small fractions of genes that did not generate good design for six encoding probes were removed, leaving 22,950 genes total, 19,717 genes in Tier 1 and the 3,233 genes in Tier 2. In Tier 1, a small number of genes had 5 probes (304 genes) and 4 probes (184 genes).

Transcripts for our isoform library were chosen from the Tier 1 and Tier 2 genes, as described above. We then performed the following additional filtering steps. First, we removed all transcripts with an FPKM less than 0.1 in bulk sequencing data.^41^ Next, we included only transcripts with Transcript Support Level (TSL) of one (as reported by Ensembl, release 111). Next, we removed all genes with only one remaining isoform. From the remaining genes, we selected those genes with at least 2 isoforms that could be uniquely targeted with 6 MERFISH probes, resulting 4,053 genes and 9,710 isoforms. These isoforms were imaged together with the ∼23,000 genes using three back-to-back MERFISH runs with three MERFISH codebooks, Codebook 1 (Tier 1 gene), Codebook 2 (Tier 2 gene), and Codebook 3 (isoforms). All MERFISH codebooks are provided in **Table S1**.

### Construction of MERFISH encoding probes

The pre-amplified encoding probe libraries were obtained from Twist Biosciences (**Table S2**). Each probe library was purchased as a separate pool. These libraries were amplified as previously described,^38^ using limited cycle PCR (Phusion Polymerase, NEB M0536L) and monitored via qPCR. The libraries were then converted to RNA via in vitro T7 transcription (HiScribe T7 Quick High Yield Kit, New England Biolabs) from a T7 promoter integrated into the PCR product. The resulting RNA products were purified using a Zeba spin desalting column (ThermoFisher, A57764) and reverse transcribed (Maxima H- Reverse Transcriptase, ThermoFisher). The RNA in the resulting RNA:DNA hybrid was degraded using alkaline hydrolysis. The ssDNA products were then purified using in house-made magnetic beads. The final library concentration was 5-20 nM/probe.

### Construction of MERFISH readout probes

Amine-modified oligonucleotide probes were purchased from IDT (Integrated DNA Technologies) and reconstituted to a concentration of 1 mM in nuclease-free water. The probes to be imaged in the 560-nm channel contained an amine group on both the 3’ and 5’ end while the probes to be imaged in the 750-nm and 650-nm channels contained an amine group on only the 5’ end of the oligonucleotide. This strategy was used because singly-labeled 560-nm probes appeared dimmer than the 750-nm and 650-nm probes. For each readout probe, 20 µL of the oligonucleotide solution was transferred into a 1.5 mL DNA low-binding microcentrifuge tube (Eppendorf, 69700) and mixed with 23 µL of 100 mM sodium bicarbonate buffer (Fisher Scientific, J60408.AP) to reach a final reaction volume of 43 µL. DBCO-SS-NHS ester (Conju-Probe LLC, CP-2024) was reconstituted by dissolving 10 mg in 100 µL anhydrous DMSO (Fisher Scientific, D12345). For each labeling reaction, 2.8 µL of the reconstituted DBCO solution was diluted in 30 µL of DMSO and thoroughly mixed. The DBCO solution was then combined with the oligonucleotide solution, vortexed gently, and incubated at room temperature for 2 hours in the dark. Then ethanol precipitation was performed to purify the DBCO-labeled DNA. Briefly, 60 µL of 1 M sodium acetate (Fisher Scientific, AM9740), 6 µL of 1 M magnesium chloride (Fisher Scientific, AM9530G), 200 µL of nuclease-free water, and 500 µL of ice-cold 100% ethanol were added to the reaction mix. Samples were incubated at -80 °C overnight and centrifuged at 15,000 × g for 30 minutes at -15 °C. The supernatant was discarded, and the pellet was washed twice with 70% ice-cold ethanol, followed by air-drying for approximately 2 hours. The purified DBCO-labeled oligonucleotides were then reconstituted in 20 µL nuclease-free water. The DBCO-modified oligo was then labeled with an azide-modified fluorescent dye (Lumiprobe, Cy3b 19330; Sulfo-Cy7 A5330; Alexa Fluor 647 16820). Each dye (1mg) was reconstituted in 100 µL of anhydrous DMSO to create a master mix stock. For each labeling reaction, 2.6 µL of dye master mix was combined with 17.4 µL DMSO in a separate tube, mixed thoroughly, and added to the tube containing 20 µL of DBCO-labeled DNA. Reactions were protected from light and incubated overnight at room temperature. Ethanol precipitation was performed to purify the dye-conjugated oligo. To each reaction mix, 30 µL of 1 M sodium acetate, 3 µL of 1 M magnesium chloride, 100 µL of nuclease-free water, and 250 µL of 100% ethanol were added. Tubes were incubated at -80 °C overnight and centrifuged at 15,000 × g for 30 minutes at -15 °C. The supernatant was discarded, and the pellets were washed twice with 70% ethanol. Pellets were air-dried for 2 hours and reconstituted in 20 µL of nuclease-free water. Except for some of the 750-nm and 650-nm readout probes, which were purchased from biosynthesis, all readout probes were synthesized using the method described here. Readout sequences can be found in **Table S2**.

### MERFISH imaging

MERFISH experiments were performed using a custom-built epifluorescence microscope as previously described.^18^ Samples were loaded into a commercial flow chamber (FCS2, Bioptechs) equipped with a 0.75-mm-thick flow gasket (DIE F18524, Bioptechs). All buffers and readout probe mixtures were delivered using a home-built automated fluidics system. Prior to imaging, samples were stained with DAPI to visualize cell nuclei and enable generation of a low-resolution mosaic for tiling. Hybridization was performed directly on the microscope using a hybridization buffer composed of 2xSSC, 10% (v/v) ethylene carbonate (Sigma), and 0.1% Triton X-100. Readout probes were used at a final concentration of 3 nM; in cases of double-labeled probes, each probe was diluted to 1.5 nM. Each hybridization round lasted 15 minutes and was followed by a wash in the same buffer. Imaging buffer comprising 5 mM 3,4-dihydroxybenzoic acid (P5630, Sigma), 50 µM trolox quinone, 1:500 recombinant protocatechuate 3,4-dioxygenase (rPCO; OYC Americas), and 5 mM NaOH (to adjust pH to 8.0) in 2xSSC was flowed into the chamber immediately before imaging.

Each imaging round captured three color channels corresponding to three readout probes (750 nm, 650 nm, and 560 nm). Fiducial beads (excited at 488 nm) were imaged at a single z-plane in each FOV to enable drift correction across imaging rounds. Imaging was performed as an 8-plane z-stack at 1.5-μm spacing. DAPI (405 nm) and total cDNA label, Alexa Fluor647 Streptavidin (Biolegend 405237), which binds to the biotinylated TSO, were only imaged in the last round for cell segmentation. Imaging for cell segmentation was performed as a 22-plane z-stack at 0.5-μm spacing.

Following each round, dyes were cleaved from the readout probes using cleavage buffer (2X SSC and 50 mM TCEP, GoldBio), incubated for 15 minutes, and washed with 2xSSC. This cycle of hybridization, imaging, and cleavage was repeated for each round. Each codebook was imaged in 30 hybridization rounds with three readout probes per round, covering 90 bits. In total, 60 rounds of imaging were conducted to capture 180 bits when imaging the ∼23,000 genes, followed by an additional round of DAPI and cDNA imaging for cell segmentation. Imaging for each experiment took ∼5 days to complete for each coronal hemisphere brain section. When imaging ∼23,000 genes plus ∼10,000 splice isoforms, 90 rounds of imaging were conducted to capture 270 bits, lasting ∼8 days per experiment.

### Bulk RNA-seq of the mouse brain

Total RNA was extracted from a third mouse brain tissue sample, with position along the anterior-posterior axis matched to the anterior slices imaged by MERFISH, using TRIzol Reagent (Thermo Fisher) followed by column-based purification with in-column DNase treatment. Briefly, 1 mL of TRIzol was added to the tissue sample and incubated for 5 minutes at room temperature to ensure complete dissociation of nucleoprotein complexes. Phase separation was initiated by adding 0.2 mL of chloroform per 1 mL of TRIzol, followed by vigorous shaking and a 2-3-minute incubation. Samples were centrifuged at 12,000 × *g* for 15 minutes at 4 °C, and the resulting aqueous phase was carefully transferred to a new tube. An equal volume of 100% ethanol was added to the aqueous phase (1:1), mixed thoroughly, and the solution was applied to a Zymo-Spin™ IC Column (Zymo Research). After centrifugation, the flow-through was discarded. Columns were then washed with 400 µL RNA Wash Buffer. For genomic DNA removal, DNase I treatment was performed in-column by applying 40 µL of freshly prepared DNase I reaction mix (5 µL DNase I enzyme and 35 µL DNA Digestion Buffer) directly onto the column matrix, followed by a 15-minute incubation at room temperature. After DNase treatment, RNA purification continued according to the manufacturer’s RNA Clean-up protocol. Columns were washed sequentially with 400 µL RNA Prep Buffer, 700 µL RNA Wash Buffer, and a final wash with 400 µL RNA Wash Buffer. To elute the RNA, 15 µL of nuclease-free water was applied directly to the column matrix and centrifuged. Eluted RNA was stored at -80 °C until further use. Library preparation, quality control, and RNA sequencing were performed by Novogene. Total RNA samples were assessed for integrity and used to construct sequencing libraries following standard Illumina protocols. Libraries were sequenced on an Illumina platform to generate 150 bp paired-end reads. Reads were aligned to the reference genome using HISAT2, and gene expression levels were quantified for downstream analysis by Novogene using their in-house pipeline.

### MERFISH Image Analysis

All MERFISH image processing was carried out using the Python package MERlin (https://github.com/ZhuangLab/MERlin), as previously described.^18^ Briefly, fiducial beads were identified in each field of view (FOV) across all imaging rounds. These fiducial positions were used to calculate x-y stage drift relative to the first round, enabling precise alignment of all FOV images throughout the experiment. After alignment, a high-pass filter was applied to each FOV’s image stack to remove background signal. Images were then subjected to four rounds of Lucy-Richardson deconvolution to sharpen RNA spots, followed by low-pass filtering to correct for small positional shifts in RNA molecule centroids between imaging rounds. RNA molecules were then identified using our previously published pixel-based decoding algorithm implemented in MERlin by aggregating adjacent pixels assigned the same barcode into putative RNA molecules.^18^ Barcodes were filtered to retain only those transcripts with an estimated misidentification rate below 5%. The misidentification rate was estimated using blank barcodes as described previously.^18^ A random subset of blank barcodes with the number equal to ∼10% of the transcript-coding barcodes were used to evaluate misidentification rate, and these barcodes were randomly selected from the blank barcodes, while ensuring even coverage across all 90 bits to avoid bias in readout usage. As compared to our previously established MERlin pipeline, a notable modification was required for decoding the long, 90-bit MERFISH images acquired in this work. Due to the computational demands of nearest neighbor calculations with these long codebooks, we accelerated the decoding step by using the GPU-accelerated cdist function from the CuPy Python package^125^, replacing the slower CPU-based approach reported earlier.

### Cell Segmentation

Cell segmentation was performed using a two-step approach. First, we applied the computational technique Dark Sectioning^128^ to the MERFISH DAPI channel to remove out-of-focus background signal that could otherwise degrade the segmentation quality. To further reduce computation load before segmentation, image volumes were first downsampled fourfold in the lateral dimensions while retaining the original axial step size of 500 nm, resulting in nearly isotropic voxels. The processed images were then segmented with Cellpose-SAM,^126^ utilizing the default SAM model with 3D mode enabled on an A100 or H100 GPU. The resulting masks were then upsampled to their original resolution and used to generate cell polygons for partitioning barcodes into individual cells.

### Registration of MERFISH Data to the Allen Mouse Brain CCF

We registered MERFISH data to the Allen Mouse Brain Common Coordinate Framework version 3 (CCFv3) using the SimpleElastix toolkit,^127^ which enables nonlinear image transformation between a “moving” image and a “fixed” template. For each slice, the PolyT channel from the MERFISH images was used as the fixed image, while a corresponding coronal slice was selected from the Nissl channel of the Allen Reference Atlas and designated as the moving image.

To enhance registration accuracy, particularly in CCF regions with limited contrast in either the PolyT or Nissl channels, we generated 3–4 additional synthetic channels for both fixed and moving images. First, we clustered cells from each brain slice at low-resolution, generating approximately 10 discrete clusters, most clusters representing major anatomical regions. For anterior slices, these clusters correspond to areas such as cortex, rostral (rostroventral) part of the Lateral septal nucleus (LSr), striatum, and fiber tracts; for posterior slices, the clusters include structures such as cortex, dentate gyrus, thalamus, hypothalamus and ventricles. These clusters were then visualized as 2D histograms and smoothed with a Gaussian blur to generate additional fixed-image channels (**Figure S1B)**. For the moving image, we used the annotation of the Allen Reference Atlas to create masks for the same major anatomical regions, and applied these masks to the Nissl image, producing matching synthetic channels (**Figure S1B**).

Registration was performed using a multi-channel approach, beginning with an affine transformation followed by a B-spline nonlinear transformation, with all channels jointly informing the alignment. The resulting transformation effectively warped the moving Nissl image to match the PolyT channel (**Figure S1C**). Using this transformation, we then mapped the cell coordinates back to CCFv3 space, achieving close correspondence with reference anatomical boundaries (**Figure S1D**). For the spatial Hotspot analysis, we used the same transformations, but rather mapped the positions of the individual RNA transcripts back to CCFv3 space.

### Hotspot analysis of spatial gene modules

Spatial gene modules were identified using the Hotspot analysis package.^42^ We began with raw barcode data that had been registered to Allen CCFv3 coordinates. Barcodes were aggregated into spatial bins of 25 × 25 μm to generate a counts matrix representing gene expression at each spatial position. This matrix was analyzed in Hotspot using the Bernoulli model and genes exhibiting significant spatial correlation were identified with a neighborhood size of 36 nearest neighbors and a false discovery rate (FDR) threshold of 0.01. Correlation coefficients were then computed for each gene pair, and spatial gene modules were identified from this correlation matrix at two different resolutions, employing a minimum module size of either 300 or 30 genes. In some samples, initial module inspection revealed modules showing artifacts, such as dust particles or isolated bright spots. To address this, genes from such artifactual modules were removed, and the pairwise correlations and modules were recalculated.

For analyses involving two adjacent anterior slices, we first identified the matching set of genes showing significant spatial correlation in both samples. Pairwise correlations for these shared genes were computed separately for each sample. The resulting correlation matrices were either analyzed separately (**Figure S2A, B**) or averaged (**Figure S2C; Figure 3**) before gene module identification.

### De novo clustering and cell-type annotation

To determine the cell types in our MERFISH data, we applied Leiden clustering analysis in three successive rounds: 1) on all cells to determine the cell class labels, 2) on all neurons to determine neuronal subtypes, and 3) on cortical excitatory neurons to determine layer and projection specific classification. Initial filtering of the data involved removing cells with fewer than 50 counts, removing genes that appeared in fewer than 10 cells, and filtering out predicted doublets using the Scrublet Python package.^129^ Next, we used the Scanpy Python package^130^ for each round of clustering. The data were preprocessed using the following pipeline: 1) total counts were normalized to 10,000 and log1p-transformed, 2) genes were subsetted to highly variable genes, which were determined by consensus from applying the ‘highly_valiable_genes’ function to each sample separately, 3) z-score scaling was performed on each sample separately, and 4) the combined samples were subjected principle component analysis (PCA) and 85, 45, and 33 PCs were retained for the 3 successive rounds, respectively. To determine the number of PCs to keep, we calculated the largest PC after randomly shuffling the genes in the cell-by-gene matrix and used the median over ten such randomization as a cutoff threshold for retaining PCs.^134^ Next, cells were embedded in a k-nearest neighbor graph and community detection was performed using the Leiden modularity optimization algorithm,^131^ using the ‘neighbors’ and ‘leiden’ function within Scanpy (with k = 25 and resolution = 2). To determine the classification labels of the cell clusters, except for hypothalamic and midbrain neurons, we utilized a combination of established marker gene identification, spatial location, and cross-validation with labels transferred from published single-cell sequencing data.^40^ To assign hypothalamic and midbrain neurons, we used spatial location based on CCF alignment.

### Integration of MERFISH data with scRNA-seq data and cell-type label transfer

We integrated MERFISH data with single-cell RNA sequencing (scRNA-seq) data from the Allen Institute.^40^ To ensure anatomical consistency, we filtered the scRNA-seq dataset to retain only cells originating from regions assayed in the MERFISH experiments. Specifically, we selected cells from the isocortex, midbrain, striatum, hippocampal formation, hypothalamus, thalamus, pallidum, olfactory area, and cortical subplate. We integrated the two datasets using the genes that are highly variable in at least one of the datasets. The highly variable genes are computed using the highly_variable_genes function from the Scanpy Python package.^130^ After subseting the genes to the highly variable genes, each dataset was preprocessed independently using the Scanpy pipeline: total counts per cell were normalized to 1,000, log1p-transformed, and scaled to Z-scores. Following preprocessing, the two datasets were concatenated and subjected to principal component analysis (PCA), retaining the top 100 components. Integration was performed using the SeuratIntegration class from the ALLCools Python package, a canonical correlation analysis (CCA)-based integration algorithm.^132, 135^ CCA was applied to co-embed the datasets in a 100-dimensional space. For each cell, we identified its five nearest neighbors across datasets, and cell pairs that were mutual nearest neighbors were designated as integration anchors. Finally, we used the label_transfer function from the SeuratIntegration class to transfer cell-type annotations (class and subclass) from the scRNA-seq dataset to the MERFISH dataset via the identified anchors. This function estimates, for each query MERFISH cell, the probability of a given label as the fraction of scRNA-seq cells with that label among its 100 nearest-neighbor anchors in PCA space. The label with the highest probability was assigned to the query cell, and this probability was reported as the label transfer confidence score.

### Spatial differential gene expression analysis

We performed differential expression (DE) analysis between the isocortex and all other brain regions for each cell type. Cells were grouped by anatomical region using the Allen CCF annotations, and DE analysis was performed separately for each cell type using the Wilcoxon rank-sum test as implemented in Scanpy’s rank_genes_groups function. All analyses were conducted on log-normalized expression values. Adjusted p-values were computed using the Benjamini-Hochberg method, and genes with a false discovery rate (FDR) < 0.05 and absolute log2 fold change > 0.25 were considered significant. Gene ontology (GO) enrichment analysis was performed using g:Profiler (via the gprofiler-official Python package)^133^, which tests for overrepresentation of functional categories based on annotated gene sets. Enrichment was conducted using the GO:BP (Biological Process), GO:MF (Molecular Function), and GO:CC (Cellular Component) ontologies with term size <2,000 genes.

To investigate transcriptional differences associated with local cell density of SMCs, we first quantified the number of same-type neighbors for each cell using spatial coordinates. For each sample, we extracted the 2D spatial positions and constructed a cKDTree to identify all neighboring cells within a 100-μm radius. For each cell, we counted the number of neighboring cells of the same cell type. Next, we classified cells as “dense” or “sparse” SMCs based on the distribution of their neighbor counts. We assigned the top 20% (highest neighbor counts) as “dense” and the bottom 20% as “sparse.” We then performed DE analysis between dense and sparse SMCs. Genes with adjusted p-values < 0.05 and absolute log2 fold change ≥ 0.25 were considered significant.

### Ligand-receptor analysis

To quantify spatial colocalization of ligand-receptor (LR) pairs, we calculated colocalization scores for all LR pairs present in our dataset using the CellTalkDB reference list.^62^ About 1,500 of ∼2,000 total pairs in the CellTalkDB reference list were present in our imaged gene panel. Analyses were performed on log-transformed and normalized expression data, with an expression threshold greater than zero to define ligand+ and receptor+ cells. For each LR pair, we identified ligand+ or receptor+ cells in each sample and extracted their spatial coordinates. Spatial proximity between ligand+ and receptor+ cells was assessed using a 100-µm radius, with a separate cKDTree constructed for each sample. For each ligand+ cell, we determined whether any receptor+ cells were present within the specified radius (ligand-to-receptor proximity), and vice versa (receptor-to-ligand proximity). Colocalization scores were computed as the total number of such proximity events (in both directions) normalized by the combined number of ligand+ and receptor+ cells.

To infer ligand-receptor interactions between cell types from proximity, we first identified all receptor+ cells and identified all ligand+ cells within a 100 µm radius of each receptor+ cell. Every receptor-ligand cell pair within this radius was recorded, producing a one-to-one interaction table that included cell IDs, spatial distances, expression levels, and cell type annotations for both the receptor+ and ligand+ cells. To focus on receptor cells with a meaningful local ligand microenvironment, we filtered out any receptor+ cell that had fewer than five ligand+ neighbor cells within the 100 µm radius. To reduce complexity, fine-grained cell type labels were grouped into broader categories. The “Cortex_IT_excitatory” neuron group included L2/3 IT, L4/5 IT, L2/3 IT RSP, L4 IT RSP, L5 IT, and L6 IT. The “Cortex_non_IT_excitatory” neuron group included L5 NP, L5 ET, L6 CT, and L6b. The “Inhibitory_neuron” group included Inhibitory_Lamp5, Inhibitory_Vip, Inhibitory_Sst, and Inhibitory_Pvalb. The “Hippocampus-neuron” group included Hippocampus_IG, Hippocampus_CA1, Hippocampus_CA2-CA3, and Hippocampus_DG. The “Striatum_neuron” group included Striatum_D1, Striatum_D2, Striatum_LSr, and Striatum_OT. The “Cholinergic_neuron” group included Cholinergic_STR-HY and Cholinergic_TH. For analysis of the Dpyn-Oprk1 pair, the “Str neuron” group was further subdivided into Str_D1 neuron, Str_D2 neuron, and Str_other, which included Striatum_LSr and Striatum_OT. Each receptor-ligand interaction was annotated with the above-described cell-type identity of the receptor+ and ligand+ cells. These annotations were used to categorize interactions by cell-type combinations (e.g., Microglia-Astrocyte). We then aggregated all individual cell-cell interactions by computing the total number of interacting (proximal) cell pairs per cell-type pair. For each pair of interacting cells, we summed the ligand and receptor expression values; then, for each cell-type pair, we calculated the average of these sums across all interacting cell pairs in that group. This produced a cell-type-level interaction matrix that captured both the frequency of interactions between each cell type pair and the associated ligand-receptor expression levels. To compare the contributions to each LR interaction from different cell-type pairs, we determined the fraction of the interacting cell-pairs for this LR pair from each cell-type pair. These data were visualized in a dot plot where dot size represented the fraction of total interaction cell-pairs belong to the cell-type pair and dot color indicated the average expression sum of ligand and receptor.

### Splice isoform analysis

To evaluate splice isoform usage patterns within each gene, we computed Shannon entropy based on the relative expression of each detected isoform. More specifically, for each gene, we summed the total barcode counts across all detected isoforms and computed the fractional abundance *pi* of each isoform *i* relative to the gene’s total expression. Shannon entropy (H) was then calculated from these fractions to capture how evenly expression was distributed across isoforms.

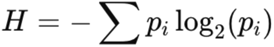

To enable comparisons across genes with differing isoform numbers, we normalized each gene’s entropy by the theoretical maximum entropy achievable for that number of isoforms,

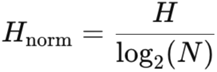

where *N* is equal to the number of isoforms for that gene. This normalized entropy metric ranges from 0 (complete dominance by one isoform) to 1 (equal usage of all isoforms), providing a continuous measure of isoform usage across genes. Shannon entropy was computed on aggregated data across three replicates.

To assess how evenly isoforms of a gene were expressed within different brain regions, we computed region-specific Shannon entropy to assess isoform usage patterns within each brain division. For each gene, barcode counts were summed across all detected isoforms within each region, and isoform fractions were calculated relative to the gene’s total regional expression within the region, as described above. Shannon entropy was then computed per gene-region pair and normalized by the maximum possible entropy for the number of isoforms targeted for that gene. This yielded a normalized entropy score, enabling comparisons of isoform diversity across anatomical divisions.

To identify genes exhibiting different spatial localization patterns among isoforms, we also performed an alternative spatial analysis of the isoforms based on the afore-mentioned Hotspot analysis using all ∼33,000 measured transcripts. We ran Hotspot at an intermediate resolution using a module size set to 100 genes. Then, considering only genes that have more than one isoforms being probed, we identified candidate genes in which at least two isoforms were assigned to different modules, suggesting that these isoforms adopt divergent spatial distribution. These candidates were subsequently validated by visual inspection.

To identify cell-type-specific differences in isoform usage, we performed pairwise Fisher’s exact tests on a total of 128,261 cells across all three replicates. For each gene under consideration, all possible pairs of isoforms were systematically tested within each annotated cell type. Within each cell type, we determined whether each isoform was detected in a given cell. For each isoform pair, we constructed a 2X2 contingency table that captured the number of cells in which each isoform was detected versus not detected. This allowed us to directly compare the detection frequencies of the two isoforms across the same population of cells. Fisher’s exact test was used to evaluate statistical significance, and Benjamini–Hochberg multiple hypothesis correction was applied to calculated the false discovery rate. Only values with adjusted p-value < 0.0005 were retained. For each isoform pair and cell type, we computed the log₂ fold change in fractions of cells expressing the isoforms: Log2(fraction of cells expressing isoform 1/ fraction of cells expressing isoform 2). For this analysis, excitatory IT neurons in the isocortex, including L2/3 IT, L4/5 IT, L5 IT, L6 IT, L2/3 IT RSP, and L4 IT RSP, were grouped together under the label Cortex_IT_excitatory neurons. Other excitatory neurons in the isocortex, including L5 ET, L5 NP, L6 CT, and L6b, and Cortex L5 NP, were grouped as Cortex_non_IT_excitatory neurons. All other cell types retained their original annotations.

## Supplementary Figures

**Figure S1:**
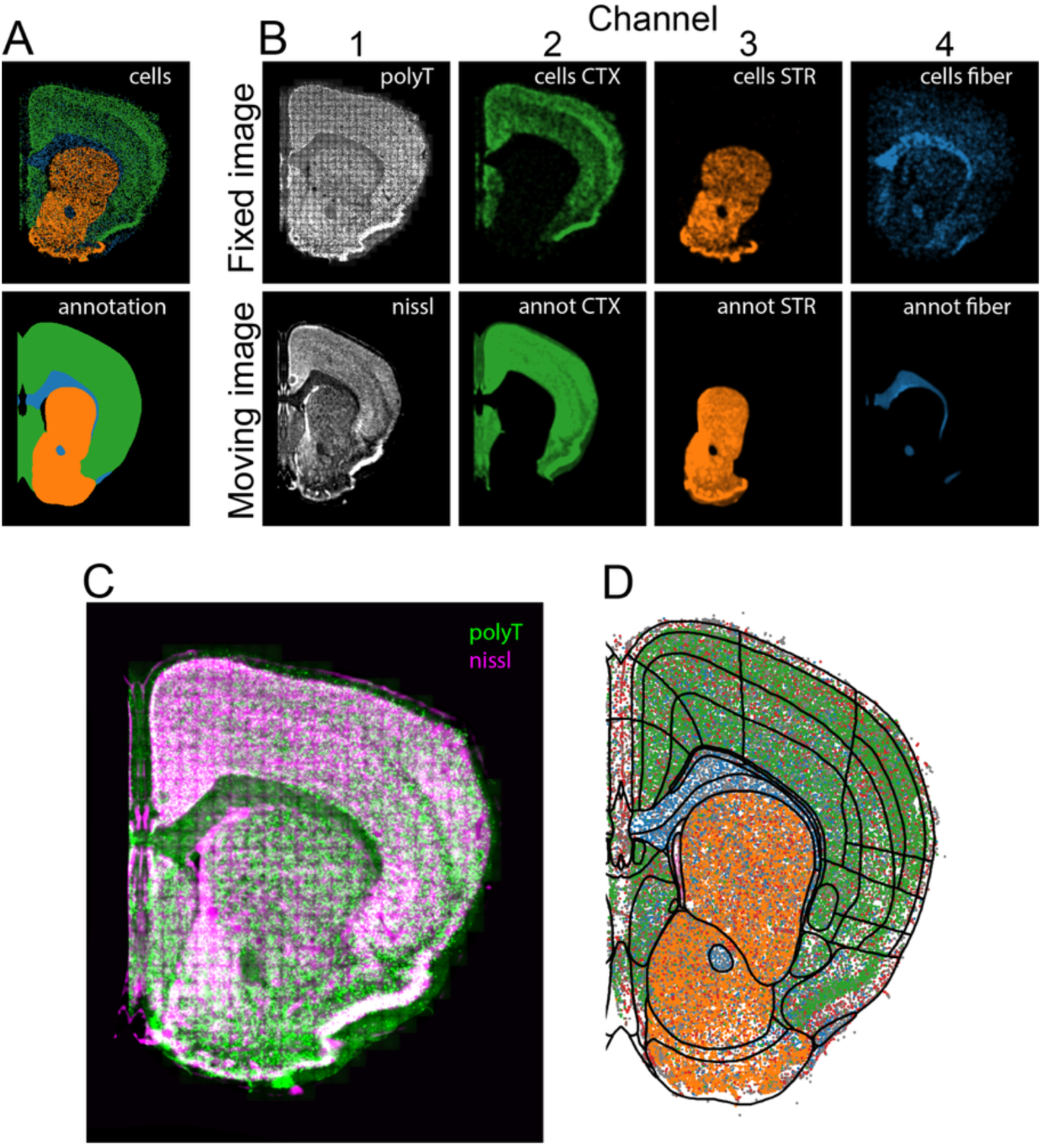
Registration of MERFISH images to the Allen CCF version 3. (A) Selected cells from low-resolution clustering localizing to the isocortex, fiber tract and striatum (top) and corresponding regions identified in the CCF annotation (bottom). (B) Individual channels of the multichannel registration. The fixed images (top) included PolyT, and images rendered from the cells identified in the three regions indicated in (A) measured in the RT&T-AMP-MERFISH experiment. The moving images (bottom) included the Allen nissl image and individual nissl images masked by the corresponding annotation regions in Allen CCF. (C) Resulting nissl image warped to the polyT image. (D) Cell positions measured in the RT&T-AMP-MERFISH experiment transformed to Allen CCF coordinates after registration. The black lines demarcate brain regions defined in the Allen CCF.

**Figure S2:**
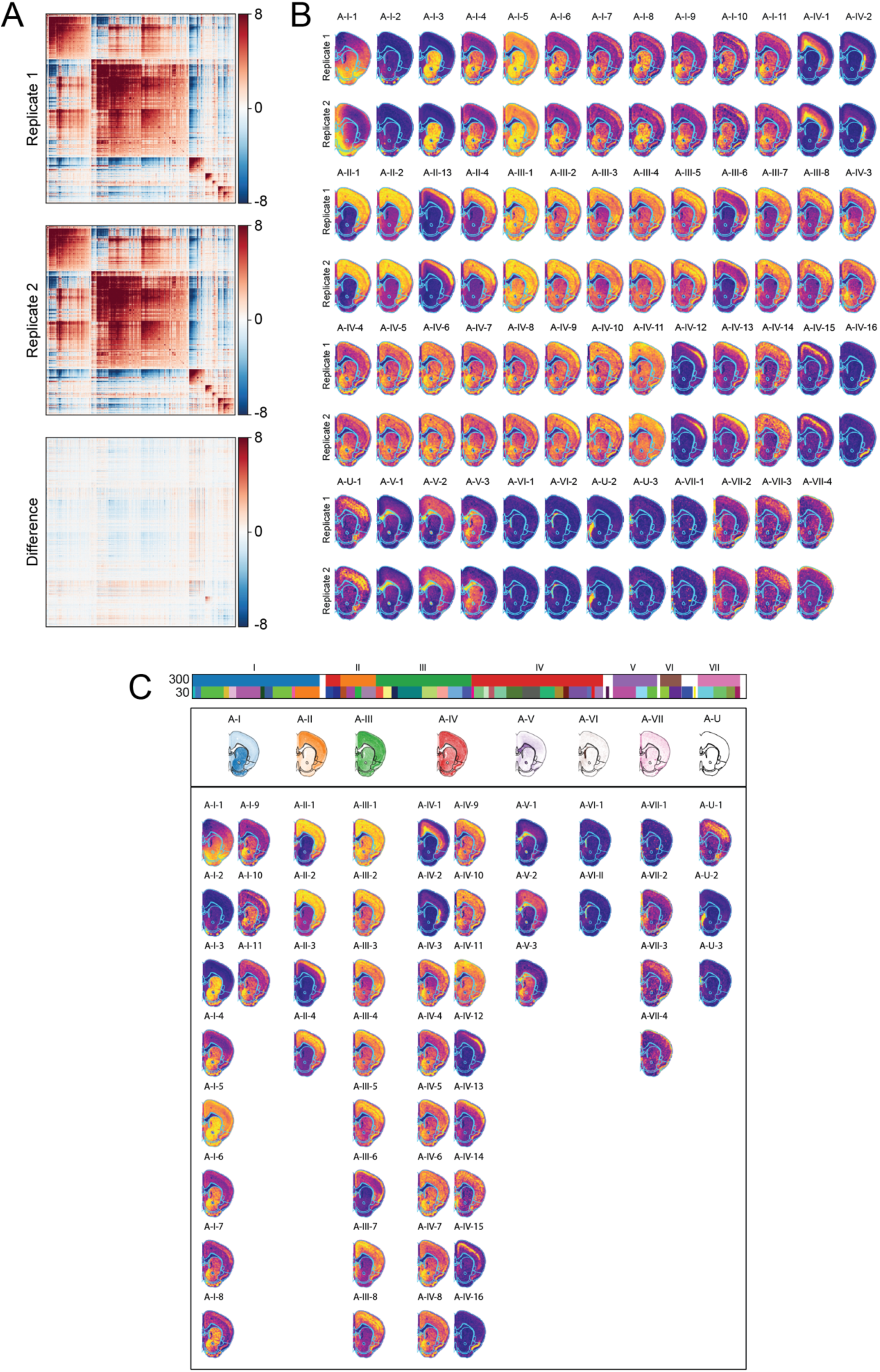
Spatial Hotspot analysis of the two adjacent anterior slices. (A) Spatial correlation matrices of two adjacent anterior slices (top and middle panels) and the difference matrix between the two slices (bottom panel). Because the two slices are adjacent to each other (10 μm apart), they serve as replicates for us to evaluated reproducibility of the Hotspot analysis. The correlation matrices are nearly identical to each other, indicating high reproducibility. (B) High-resolution modules from these two anterior slices. Among the 51 spatial modules, all but one showed highly similar spatial patterns, indicating high reproducibility. The distinct one (A-I-1) does not align to any anatomic structure in the brain and likely results from an artifact. (C) Average Hotspot analysis of the two anterior slices. Because the two adjacent slices showed nearly identical Hotspot analysis results, we averaged the correlation matrix and identified the spatial modules from the average matrix. The spatial maps of the modules are shown in a hierarchical view with the low-resolution modules (minimum 300 genes) denoted with Roman numerals (I, II, III, …) and high-resolution modules (minimum 30 genes) denoted with Arabic numerals (1, 2, 3…). “A” in the module annotation stands for “anterior”.

**Figure S3:**
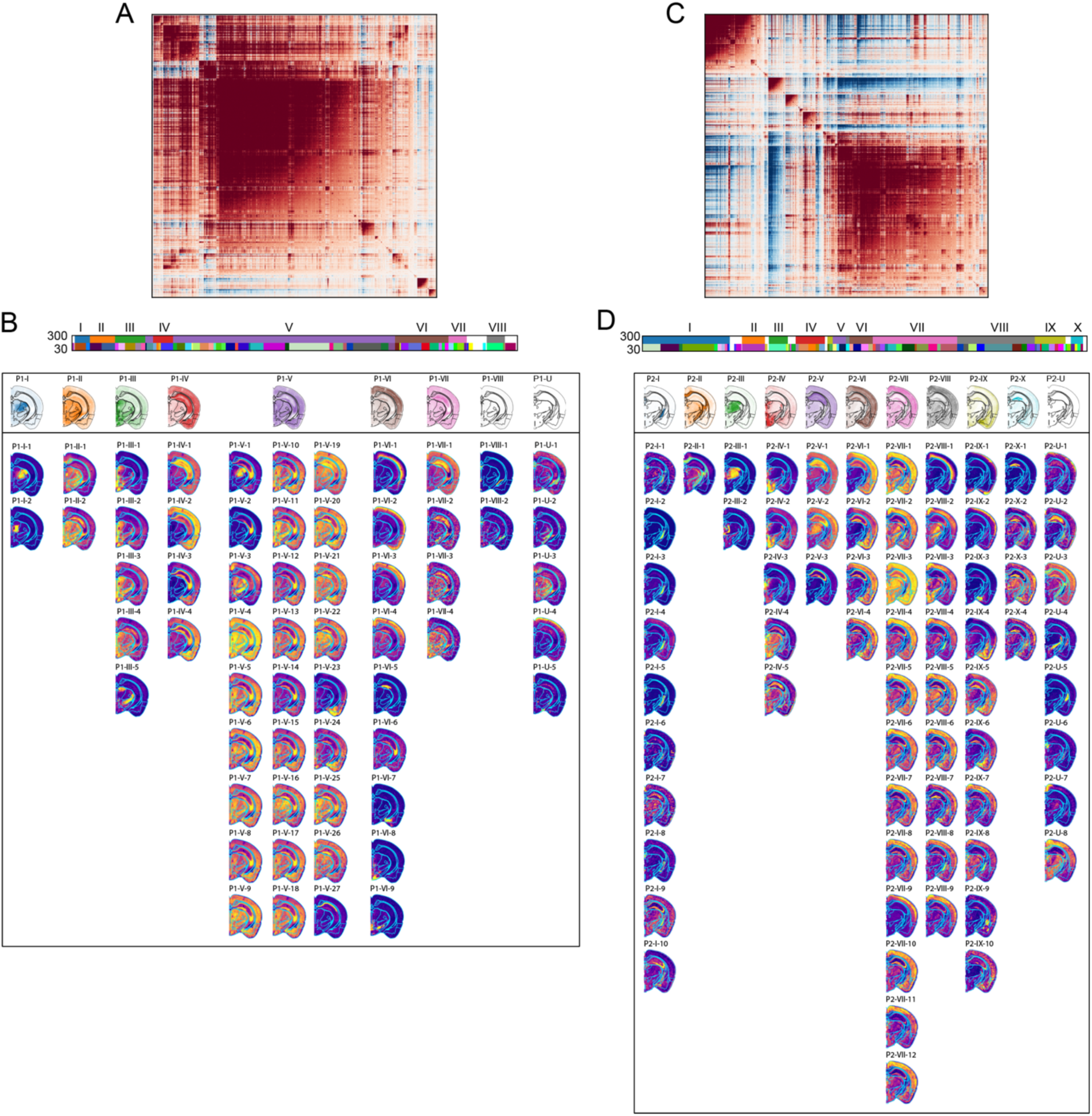
Spatial hotspot analysis of the two posterior slices. (A-D) Spatial Hotspot analysis results for two posterior slices shown separately, with modules named and displayed as in Figure S2. (A) and (B) are the correlation matrix and spatial maps of gene modules for posterior slice 1. (C) and (D) are for posterior slice 2. “P1” and “P2” in the module names stand for posterior slices 1 and 2, respectively. Because the two posterior slices are further apart and anatomic structures in this region show relatively high-frequency variations along the anterior-posterior axis, Hotspots analysis generated partially overlapping but distinct results.

**Figure S4:**
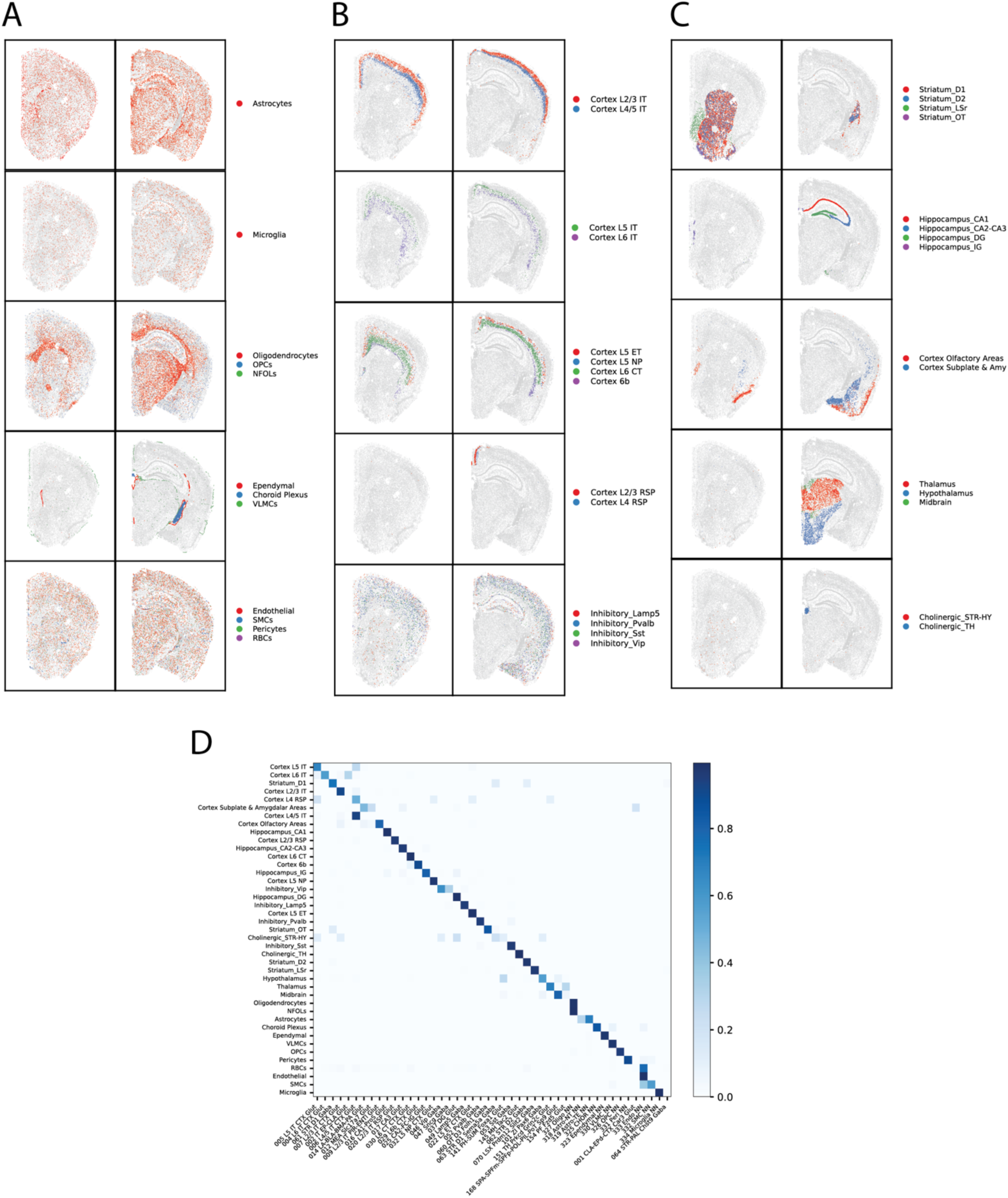
Spatial maps of major cell classes and neuronal subtypes derived from RT&T-AMP-MERFISH and the confusion matrix between *de novo* clustering results from RT&T-AMP-MERFISH data and label-transfer results from scRNA-seq data. (A) Spatial maps of non-neuronal cell classes as shown in Figure 4A, in one representative anterior slice and one representative posterior slice. (B) Spatial maps of cortical neuronal subtypes as shown in Figure 4B, in the anterior and posterior slices as in (A). (C) Spatial maps of non-cortical neuronal subtypes as shown in Figure 4B, in the anterior and posterior slices as in (A, B) (D) Confusion matrix showing the correspondence between the *de novo* cell type classification results from RT&T-AMP-MERFISH data, shown in Figure 4A**, B**, and the cell-type classification results determined by label-transfer from previous cell-type annotation based on scRNAseq data,^40^ obtained by integrating our RT&T-AMP-MERFISH data with the scRNA-seq data. Color indicates the fraction of cells in each *de novo* cluster that shares each label-transferred cell-type label. Only cells with a label transfer confidence ratio greater than 0.5 are included (**Methods**). For the label-transferred cell types in the x -axis, only cell types containing >200 cells are shown.

**Figure S5:**
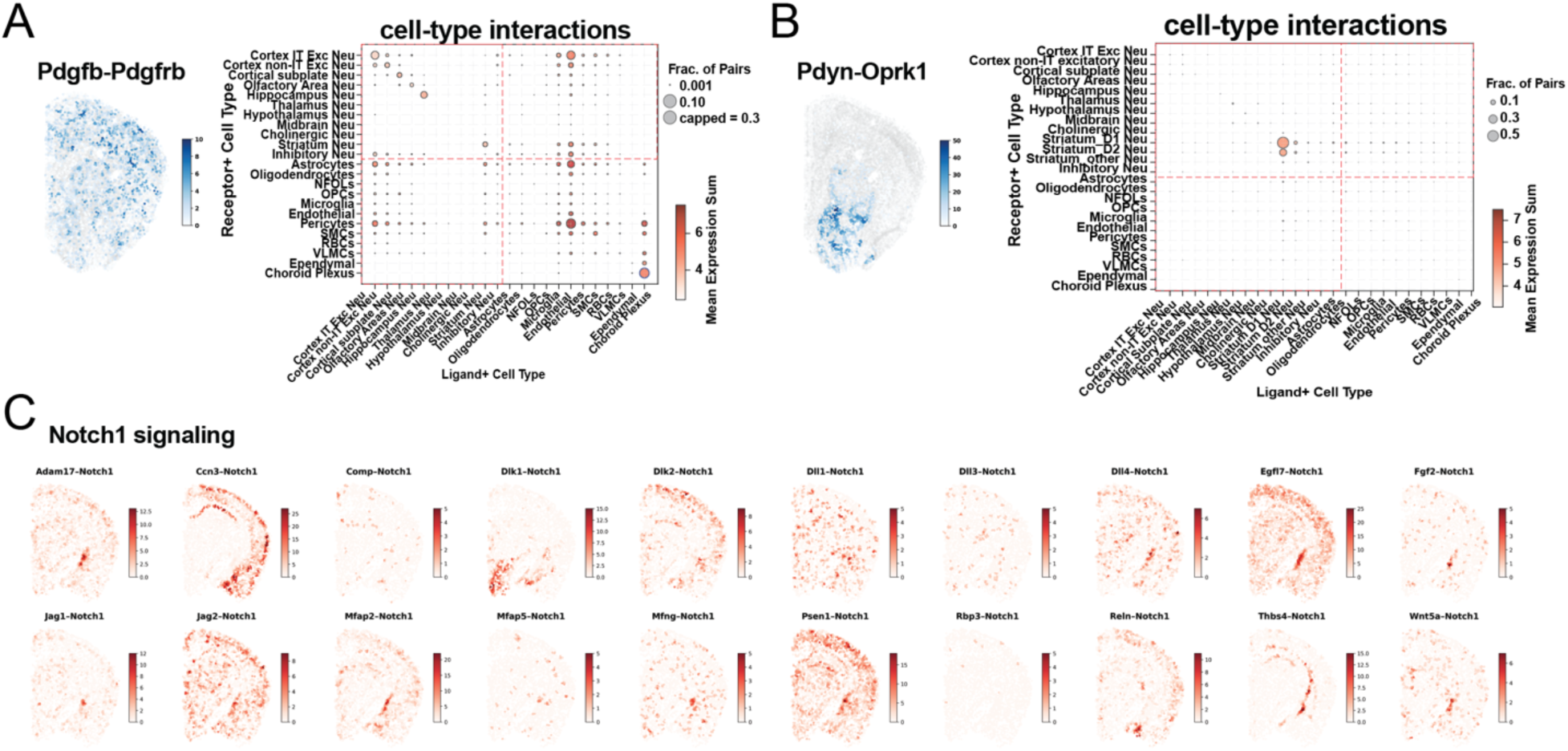
Additional ligand–receptor interaction analysis. (A) Spatial and cell-type distributions of the colocalized *Pdgfb*–*Pdgfrb* ligand–receptor pairs. Left: Spatial maps show, for each receptor-positive cell, the number of ligand-positive cells within a 100 µm radius in a representative anterior slice. Right: Dot plots indicate the sum of ligand and receptor expression (dot color) and the fraction of total interacting cell pairs for this LR pair that belongs to each cell-type pair (dot size). This latter metric evaluates the contribution of each cell-type pair to the observed LR interaction. (B) As in (A) but for the Pdyn–Oprk1 pair. (C) As in Figure 4F, but for a representative posterior brain slice.

**Figure S6:**
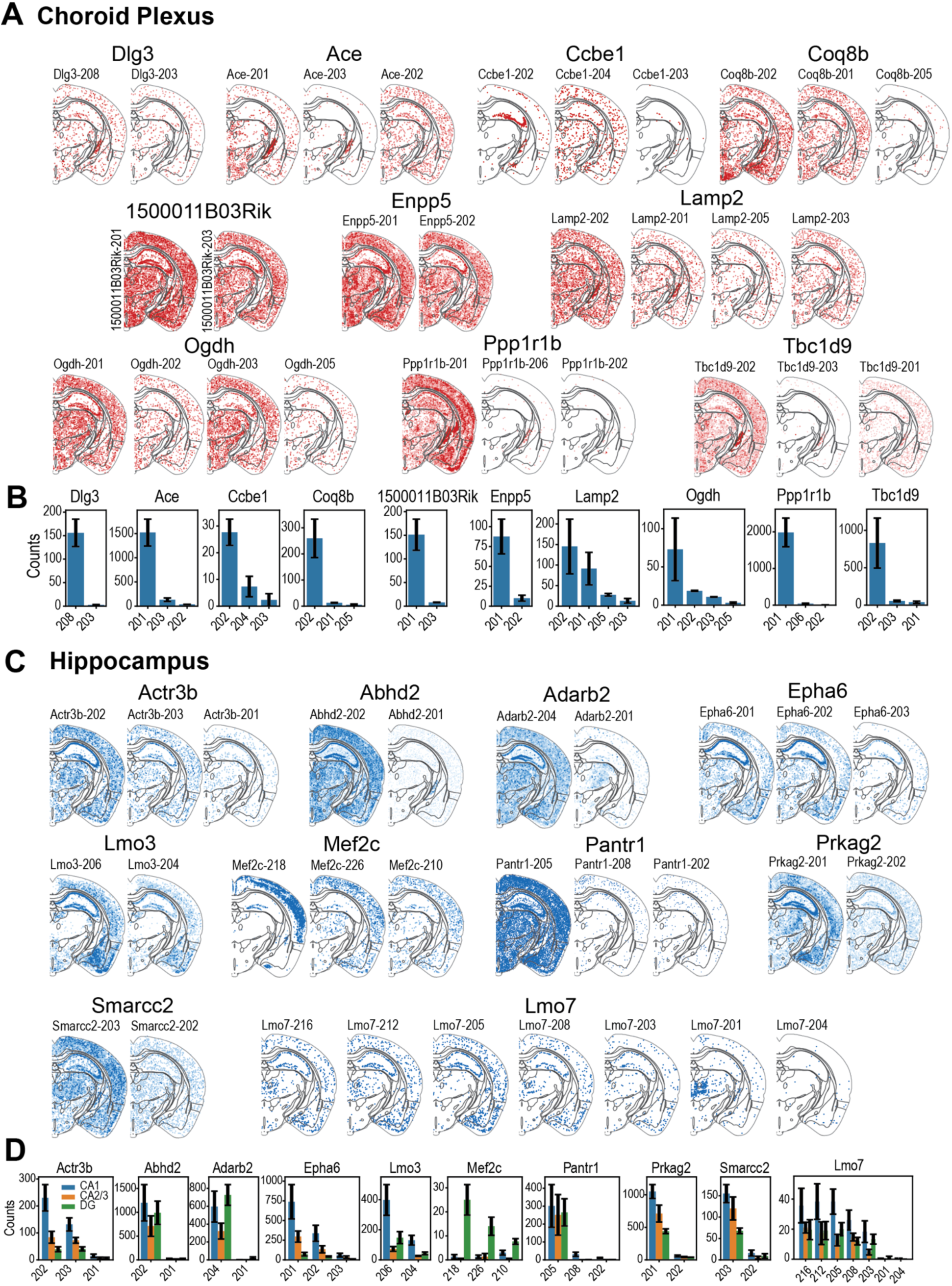
Additional examples of genes with strong specificity for isoform usage in the CP and hippocampus. (A) Spatial maps of additional example genes exhibiting strong specificity for isoform usage in the CP. (B) Expression levels (in mean ± SEM of counts, across replicates) in CP cells for the different isoforms of the genes shown in (A). (C) Spatial maps of additional example genes exhibiting strong specificity for isoform usage in the hippocampus or subregions (CA1, CA2/3 and DG) of the hippocampus. (D) Expression levels (in mean ± SEM of counts, across replicates) in CA1, CA2/3, and DG cells for the different isoforms of genes shown in (C).

## Supplementary Table Captions

**Table S1: MERFISH codebooks for the ∼23,000 genes and ∼10,000 isoforms imaged in this work.**

Three MERFISH codebooks used in the experiments indicating the 90-bit barcodes assigned to each of the genes and blank barcode controls. For the two whole-transcriptome codebooks (codebook 1 and codebook 2), the first column lists the gene name. For the isoform codebook (codebook 3), the first column lists the Ensembl transcript ID, and the second column lists gene name. The subsequent 90 columns represent the binary barcode assigned to each target, with each column named according to its corresponding readout sequence. Barcodes designated as blank controls are labeled with a gene name prefixed by “Blank-.”

**Table S2: Encoding probe, readout probe, primer sequences for amplifying encoding probes, smFISH probes, and oligos used in RT&T-AMP.**

For encoding probes used for genes in codebook 1 and codebook 2, the encoding probe sequences and the target genes are indicated. For encoding probes used for isoforms in codebook 3, the encoding probe sequences, target genes, and transcript ID for the isoforms are indicated. For the readout probes, the readout probe sequence names, the color channels (in nm for wavelength) used to detect the attached dyes, and the readout probe sequences are indicated. From the PCR primers used to amplify each encoding probe library, the codebook names and primer sequences are listed. For smFISH targeting the *Sst* transcript, the sheet includes the following: (i) Probe sets used for detection of *Sst* in single-color amplified and unamplified smFISH, consisting of 30, 5, or 1 probe(s) per set, as well as readout probes and associated fluorophore color (**Figure 1B**); (ii) Probe sets used to assess signal colocalization in amplified two-color smFISH for *Sst*, consisting of 5 or 1 probe(s) per set, as well as sequences for 5 scrambled, non-targeting control probes for evaluating nonspecific signal (**Figure 1C**). Readout probes and associated fluorophore colors these probes are also included. “750” means 750 nm; “650” means 750 nm. For the oligos used in RT&T-AMP, the reverse transcription (RT) primer and template switching oligo (TSO) are listed

**Table S3: Hotspot modules.**

This table lists the gene and module correspondence for the anterior slices and for the posterior slices 1 and 2. For the two adjacent anterior slices, the listed modules were determined from the averaged correlation matrix of both slices. For the two posterior slices (P1 and P2), analyses were performed separately. In the module names, “A” denotes anterior, while “P1” and “P2” denote posterior slices 1 and 2, respectively. Low-resolution modules (minimum 300 genes) are labeled with Roman numerals (I, II, III, …); high-resolution modules (minimum 30 genes) are labeled with Arabic numerals (1, 2, 3, …). A value of ‘-1’ indicates that the gene was not assigned to any module.

**Table S4: Spatial DE genes: isocortex vs all other imaged regions for individual cell types.**

This table reports differential gene expression analyses comparing cells from the Isocortex region to those from non-Isocortex regions within each cell type. Each sheet corresponds to a single cell type. Only cell types with at least 50 cells in both the isocortex and non-isocortex groups were included. For each gene, the table lists the gene name (names), the Wilcoxon test statistic (scores), the log₂ fold change in expression between Isocortex and non-Isocortex cells (logfoldchanges), the raw p-value (pvals), and the adjusted p-value after multiple testing correction (pvals_adj). Positive log₂ fold change values indicate genes upregulated in the isocortex, while negative values indicate downregulation. All tested genes are listed regardless of statistical significance.

**Table S5: Ligand-receptor colocalization scores.**

This table summarizes the spatial colocalization between ligand- and receptor-expressing cells across the imaged mouse brain sections, for 1,455 ligand–receptor pairs analyzed in this work. Each row shows the ligand name, the receptor name, the LR pair (pathway), the number of ligand+ cells near receptor+ cells, within a 100 µm radius (total l_to_r), number of receptor+ cells near ligand+ cells, within a 100 µm radius (total r_to_l), the total number of ligand+ cells, the total number of receptor+ cells, and the ligand-receptor colocalization score, which is calculated by summing the total l_to_r and total r_to_l columns and dividing by the sum of the total_ligand_cells and total_receptor_cells columns.

